# Movies reveal the fine-grained organization of infant visual cortex

**DOI:** 10.1101/2023.08.22.554318

**Authors:** C. T. Ellis, T. S. Yates, M. J. Arcaro, N. B. Turk-Browne

## Abstract

Studying infant minds with movies is a promising way to increase engagement relative to traditional tasks. However, the spatial specificity and functional significance of movie-evoked activity in infants remains unclear. Here we investigated what movies can reveal about the organization of the infant visual system. We collected fMRI data from 15 awake infants and toddlers aged 5–23 months who attentively watched a movie. The activity evoked by the movie reflected the functional profile of visual areas. Namely, homotopic areas from the two hemispheres responded similarly to the movie, whereas distinct areas responded dissimilarly, especially across dorsal and ventral visual cortex. Moreover, visual maps that typically require time-intensive and complicated retinotopic mapping could be predicted, albeit imprecisely, from movie-evoked activity in both data-driven analyses (i.e., independent component analysis) at the individual level and by using functional alignment into a common low-dimensional embedding to generalize across participants. These results suggest that the infant visual system is already structured to process dynamic, naturalistic information and that fine-grained cortical organization can be discovered from movie data.

---

Studying the function and organization of the youngest human brains remains a challenge. Despite the recent growth in infant fMRI^?,1–5^, one of the most important obstacles facing this research is that infants are unable to maintain focus for long periods of time and struggle to complete traditional cognitive tasks^6^. Movies can be a useful tool for studying the developing mind^7^, as has been shown in older children^8–10^. The dynamic, continuous, and content-rich nature of movie stimuli^11, 12^ make them effective at capturing infant attention^13, 14^. Here, we examine what can be revealed about the functional organization of the infant brain during movie-watching.

We focus on visual cortex because its organization at multiple spatial scales is well understood from traditional, task-based fMRI. The mammalian visual cortex is divided into multiple areas with partially distinct functional roles between areas^15–17^. Within visual areas, there are orderly, topographic representations, or maps, of visual space^18, 19^. These maps capture information about the location and spatial extent of visual stimuli with respect to fixation. Thus, maps reflect sensitivity to polar angle, measured via alternations between horizontal and vertical meridians that define area boundaries^20, 21^, and sensitivity to spatial frequency, reflected in gradients of sensitivity to high and low spatial frequencies from foveal to peripheral vision, respectively^22^. Previously, we reported that these maps could be revealed by a retinotopy task in infants as young as 5 months of age^23^. However, it remains unclear whether these maps are evoked by more naturalistic task-designs.

The primary goal of the current study is to investigate whether movie-watching data re-capitulates the organization of visual cortex. Movies drive strong and naturalistic responses in sensory regions while minimizing task demands^11, 12, 24^ and thus are a proxy for typical experience. In adults, movies and resting-state data have been used to characterize the visual cortex in a data-driven fashion^25–27^. Movies have been useful in awake infant fMRI for studying event segmentation^28^, functional alignment^29^, and brain networks^30^. However, this past work did not address the granularity and specificity of cortical organization that movies evoke. For example, movies evoke similar activity across infants in anatomically aligned visual areas^28^, but it remains unclear whether responses to movie content differ *between* visual areas (e.g., is there more similarity of function within visual areas than between^31^). Moreover, it is unknown whether structure *within* visual areas, namely visual maps, contributes substantially to visual evoked activity. Additionally, we wish to test whether methods for functional alignment can be used with infants. Functional alignment finds a mapping between participants using functional activity – rather than anatomy – and in adults can improve signal-to-noise, enhance across participant prediction, and enable unique analyses^27, 32–34^.

Nonetheless, there are several reasons for skepticism that movies could evoke detailed, retinotopic organization: Movies may not fully sample the stimulus parameters (e.g., spatial frequencies) or visual functions needed to find topographic maps and areas in visual cortex. Even if movies contain the necessary visual properties, they may unfold at a faster rate than can be detected by fMRI. Additionally, naturalistic stimuli may not drive visual responses as robustly as experimenter-defined stimuli that are designed for retinotopic mapping with discrete onsets and high contrast. Finally, the complexity of movie stimuli may result in variable attention between participants, impeding discovery of reliable visual structure across individuals. If movies do show the fine-grained organization of the infant visual cortex, this suggests that this structure (e.g., visual maps) scaffolds the processing of ongoing visual information.

We conducted several analyses to probe different kinds of visual granularity in infant movie-watching fMRI data. First, we asked whether distinct areas of the infant visual cortex have different functional profiles. Second, we asked whether the topographic organization of visual areas can be recovered within participants. Third, we asked whether this within-area organization is aligned across participants. These three analyses assess key indicators of the mature visual system: functional specialization between areas, organization within areas, and consistency between individuals.

## Results

We performed fMRI in awake, behaving infants and toddlers using a protocol described previously^6^. The dataset consisted of 15 sessions of infant participants (4.8–23.1 months old) who had both movie-watching data and retinotopic mapping data collected in the same session (Table S1). All available movies from each session were included (Table S2), with an average duration of 540.7s (range: 186–1116s).

The retinotopic-mapping data from the same infants^23^ allowed us to generate infant-specific meridian maps (horizontal versus vertical stimulation) and spatial frequency maps (high versus low stimulation). The meridian maps were used to define regions of interest (ROIs) for visual areas V1, V2, V3, V4, and V3A/B.

As a proof of concept that the analyses we use with infants can identify fine-grained visual organization, we ran the main analyses on an adult sample. These adults (8 participants) had both retinotopic mapping data and movie-watching data. Figures S1, S2, S3, and S4 demonstrate that applying these analyses to adult movie data reveals similar structure to what we find in infants.

### Evidence of area organization with homotopic similarity

To determine what movies can reveal about the organization of areas in visual cortex, we compared activity across left and right hemispheres. Although these analyses cannot define visual maps, they test whether visual areas have different functional signatures. Namely, we correlated timecourses of movie-related BOLD activity between retinotopically defined, participant-specific ROIs (7.3 regions per participant per hemisphere, range: 6–8)^31, 35, 36^. Higher correlations between the same (i.e., homotopic) areas than different areas indicates differentiation of function between areas. Moreover, other than V1, homotopic visual areas are anatomically separated across the hemispheres, so similar responses are unlikely to be attributable to spatial autocorrelation.

Homotopic areas (e.g., left ventral V1 and right ventral V1; diagonal of Figure 1A) were highly correlated (Mean [M]=0.88, range of area means: 0.85–0.90), and more correlated than non-homotopic areas, such as the same visual area across streams (e.g., left ventral V1 and right dorsal V1; Figure 1B; Δ*_Fisher Z_* M=0.42, p*<*0.001). To clarify, we use the term ‘stream’ to liberally distinguish visual regions that are *more* dorsal or *more* ventral, as opposed to the functional definition used in reference to the ‘what’ and ‘where’ streams^17^. We found no evidence that the vari-ability in movie duration per participant correlated with this difference (r=0.08, p=.700). Within stream (Figure 1C), homotopic areas were more correlated than adjacent areas in the visual hier-archy (e.g., left ventral V1 and right ventral V2; Δ*_Fisher Z_*M=0.09, p*<*0.001), and adjacent areas were more correlated than distal areas (e.g., left ventral V1 and right ventral V4; Δ*_Fisher Z_* M=0.20, p*<*0.001). There was no correlation between movie duration and effect (Same *>* Adjacent: r=-0.01, p=.965, Adjacent *>* Distal: r=-0.09, p=.740). Additionally, if we control for motion in the correlation between areas — in case motion transients drive consistent activity across areas — then the effects described here are negligibly different (Figure S5). Hence, movies elicit distinct processing dynamics across areas of infant visual cortex defined independently using retinotopic mapping.

**Figure 1:**
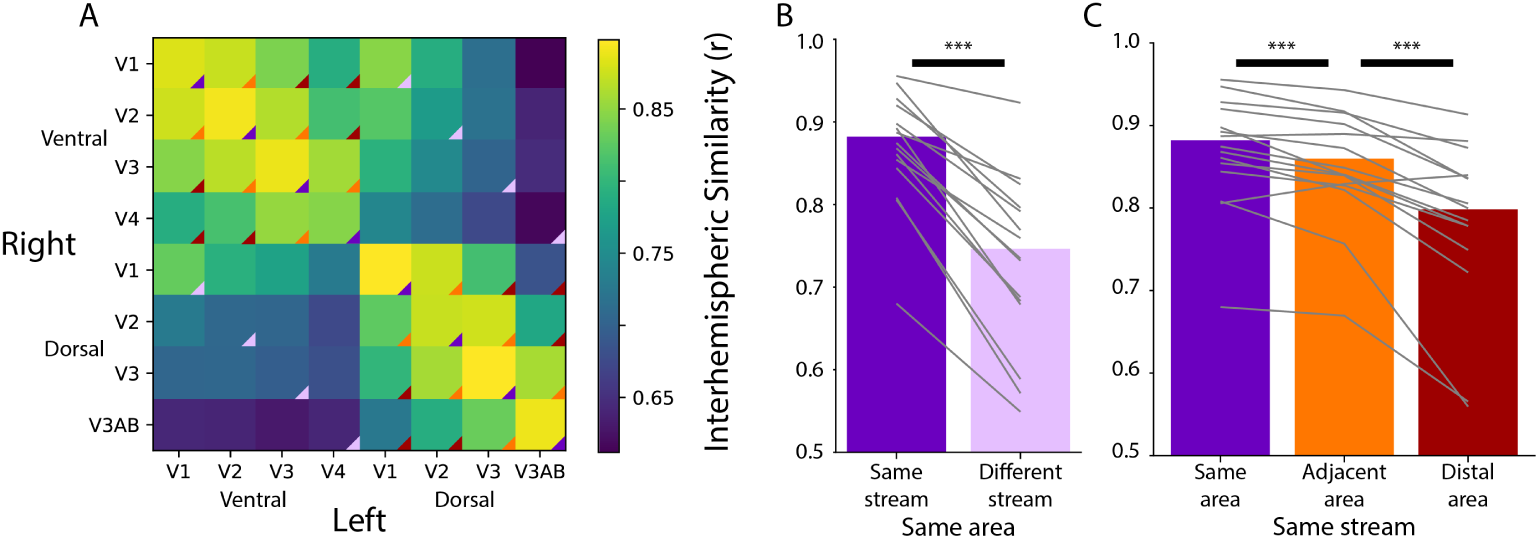
Homotopic correlations between retinotopic areas. (A) Average correlation of the time-course of activity evoked during movie watching for all areas. This is done for the left and right hemisphere separately, creating a matrix that is not diagonally symmetric. The color triangles overlaid on the corners of the matrix cells indicate which cells contributed to the summary data of different comparisons in subpanels B and C. (B) Across-hemisphere similarity of the same visual area from the same stream (e.g., left ventral V1 and right ventral V1) and from different streams (e.g., left ventral V1 and right dorsal V1). (C) Across-hemisphere similarity in the same stream when matching the same area (e.g., left ventral V1 and right ventral V1), matching to an adjacent area (e.g., left ventral V1 and right ventral V2), or matching to a distal area (e.g., left ventral V1 and right ventral V4). Grey lines represent individual participants. *** = p*<*0.001 from bootstrap resampling

We previously found^23^ that an anatomical segmentation of visual cortex^37^ could identify these same areas reasonably well. Indeed, the results above were replicated when using visual areas defined anatomically (Figure S6). However, a key advantage of anatomical segmentation is that it can define visual areas not mapped by a functional retinotopy task. This could help address limitations of the analyses above, namely that there was a variable number of retinotopic areas identified across infants and these areas covered only part of visually responsive cortex. Focusing on broader areas that include portions of the ventral and dorsal stream in the adult visual cortex^17, 37^, we tested for functional differentiation of these streams in infants. We applied multi-dimensional scaling (MDS) — a data-driven method for assessing the clustering of data — to the average cross-correlation matrix across participants (Figure S6)^35, 38^. The stress of fitting these data with a two-dimensional MDS was in the acceptable range (0.076). Clear organization was present (Figure 2): areas in the adult-defined ventral stream (e.g., VO, PHC) differentiated from areas in the adult-defined dorsal stream (e.g., V3A/B). Indeed, we see a slight separation between canonical dorsal areas and the recently defined lateral pathway^39^ (e.g., LO1, hMT), although more evidence is needed to substantiate this distinction. This separation between streams is striking when considering that it happens *despite* differences in visual field representations across areas: while dorsal V1 and ventral V1 represent the lower and upper visual field, respectively, V3A/B and hV4 both have full visual field maps. These visual field representations can be detected in adults^40^; however, they are often not the primary driver of function^38^. We see that in infants too: hV4 and V3A/B represent the same visual space yet have distinct functional profiles. Again, this organization cannot be attributed to mere spatial autocorrelation within stream because analyses were conducted across hemispheres (at significant anatomical distance) and this pattern is preserved when accounting for motion (Figure S5). These results thus provide evidence of a dissociation in the functional profile of anatomically defined ventral and dorsal streams during infant movie-watching.

**Figure 2:**
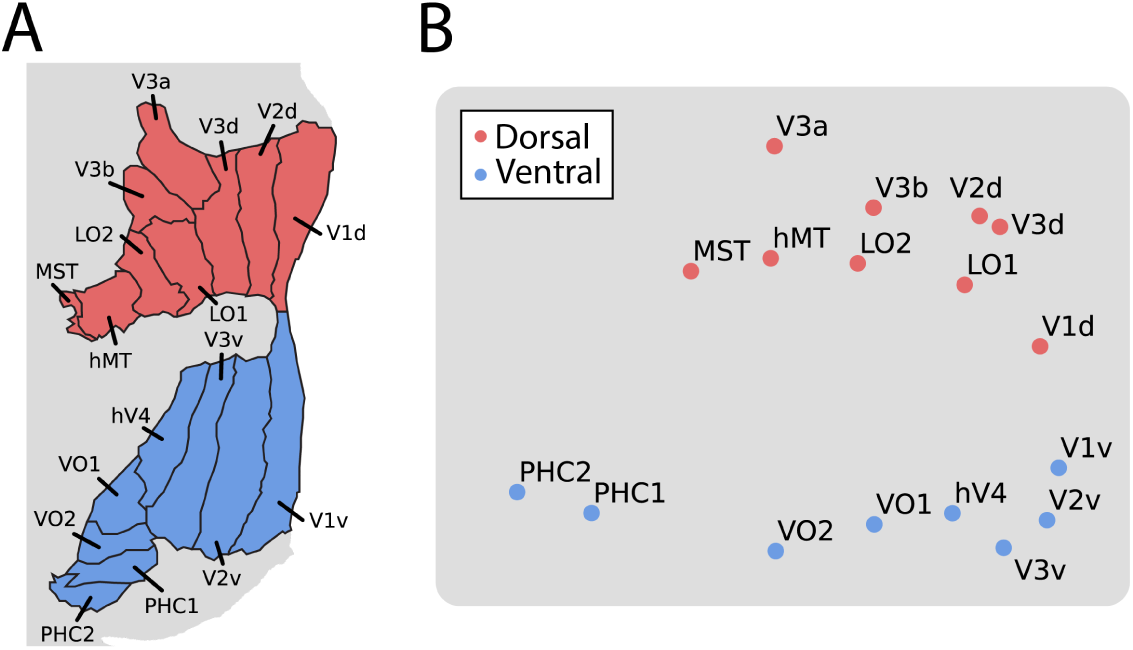
Multi-dimensional scaling (MDS) of movie-evoked activity in visual cortex. A) Anatomically defined areas^37^ used for this analysis, separated into dorsal (red) and ventral (blue) visual cortex, overlaid on a flatmap of visual cortex. B) The timecourse of functional activity for each area was extracted and compared across hemispheres (e.g., left V1 was correlated with right V1). This matrix was averaged across participants and used to create a Euclidean dissimilarity matrix. MDS captured the structure of this matrix in two dimensions with suitably low stress. The plot shows a projection that emphasizes the similarity to the brain’s organization.

### Evidence of within-area organization with independent component analysis

We next explored whether movies can reveal fine-grained organization *within* visual areas by using independent component analysis (ICA) to propose visual maps in individual infant brains^25, 26, 35, 41, 42^. ICA is a method for decomposing a source into constituent signals by finding components that account for independent variance. When applied to fMRI data (using MELODIC in FSL), these components have spatial structure that varies in strength over time. Many of these components reflect noise (e.g., motion, breathing) or task-related signals (e.g., face responses), while other components reflect the functional architecture of the brain (e.g., topographic maps)^25, 26, 35, 41, 42^. We visually inspected each component and categorized it as a potential spatial frequency map, a potential meridian map, or neither. This process was blind to the ground truth of what the visual maps look like for that participant from the retinotopic mapping task, simulating what would be possible if retinotopy data from the participants were unavailable. Success in this process requires that 1) retinotopic organization accounts for sufficient variance in visual activity to be identified by ICA and 2) experimenters can accurately identify these components.

Multiple maps could be identified per participant because there were more than one candidate that the experimenter thought was a suitable map. Across infant participants, we identified an average of 2.4 (range: 0–5) components as potential spatial frequency maps and 1.1 (range: 0–4) components as potential meridian maps. To evaluate the quality of these maps, we compared them to the ground truth of that participant’s task-evoked maps (Figure 3). Spatial frequency and meridian maps are defined by their systematic gradients of intensity across the cortical surface^43^. Lines drawn parallel to area boundaries show monotonic gradients on spatial frequency maps, with stronger responses to high spatial frequency at the fovea, and stronger responses to low spatial frequencies in the periphery (Figure S7). By contrast, lines drawn perpendicular to the area boundaries show oscillations in sensitivity to horizontal and vertical meridians on meridian maps (Figure S8). Using the same manually traced lines from the retinotopy task, we measured the intensity gradients in each component from the movie-watching data. We can then use the gradients of intensity in the retinotopy task-defined maps as a benchmark for comparison with the ICA-derived maps.

**Figure 3:**
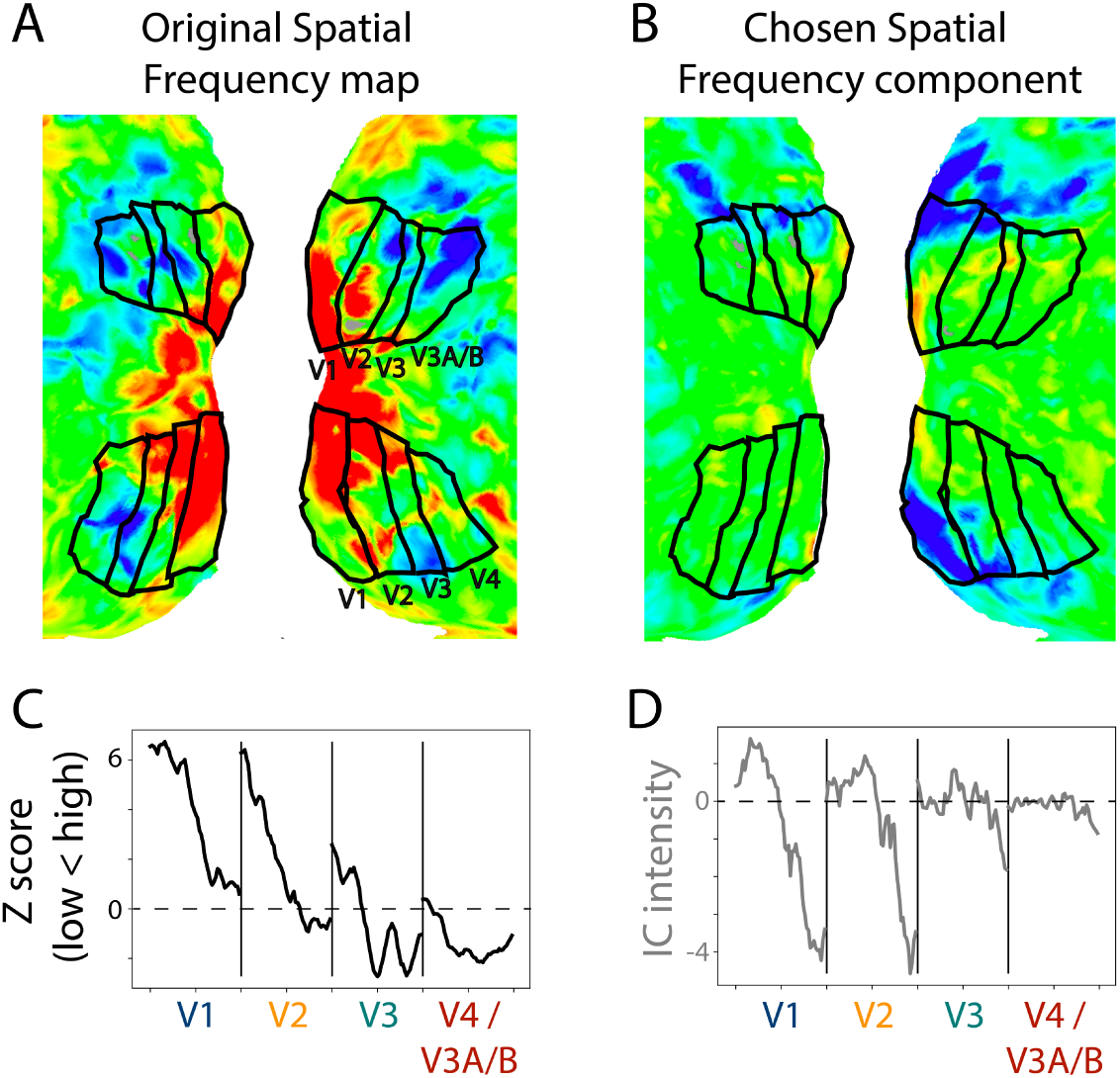
Example retinotopic task vs. ICA-based spatial frequency maps. A) Spatial frequency map of a 17.1 month old toddler. The retinotopic task data are from a prior study^23^. The view is of the flattened occipital cortex with visual areas traced in black. B) Component captured by ICA of movie data from the same participant. This component was chosen as a spatial frequency map in this participant. The sign of ICA is arbitrary so it was flipped here for visualization. C) Gradients in spatial frequency within-area from the task-evoked map in subpanel A. Lines parallel to the area boundaries (emanating from fovea to periphery) were manually traced and used to capture the changes in response to high versus low spatial frequency stimulation. D) Gradients in the component map. We used the same lines that were manually traced on the task-evoked map to assess the change in the component’s response. We found a monotonic trend within area from medial to lateral, just like we see in the ground truth. This is one example result, find all participants in Figure S7.

To assess the selected component maps, we correlated the gradients (described above) of the task-evoked and component maps. This test uses independent data: the components were defined based on movie data and validated against task-evoked retinotopic maps. Figure 4A shows the absolute correlations between the task-evoked maps and the manually identified spatial frequency components (M=0.52, range: 0.23–0.85). To evaluate whether movies are a viable method for defining retinotopic maps, we tested whether the task-evoked retinotopic maps were more similar to manually identified components than other components. We identified the best component in 6 of 13 participants (Figure 4B). The percentile of the average manually identified component was high (M=63.8 percentile, range: 26.7–98.1) and significantly above chance (ΔM=13.8, CI=[3.3–24.0], p=.011). This illustrates that the manually identified components derived from movie-watching data are similar to the spatial frequency maps derived from retinotopic mapping. The fact that this can work also indicates the underlying architecture of the infant visual system influences how movies are processed.

**Figure 4:**
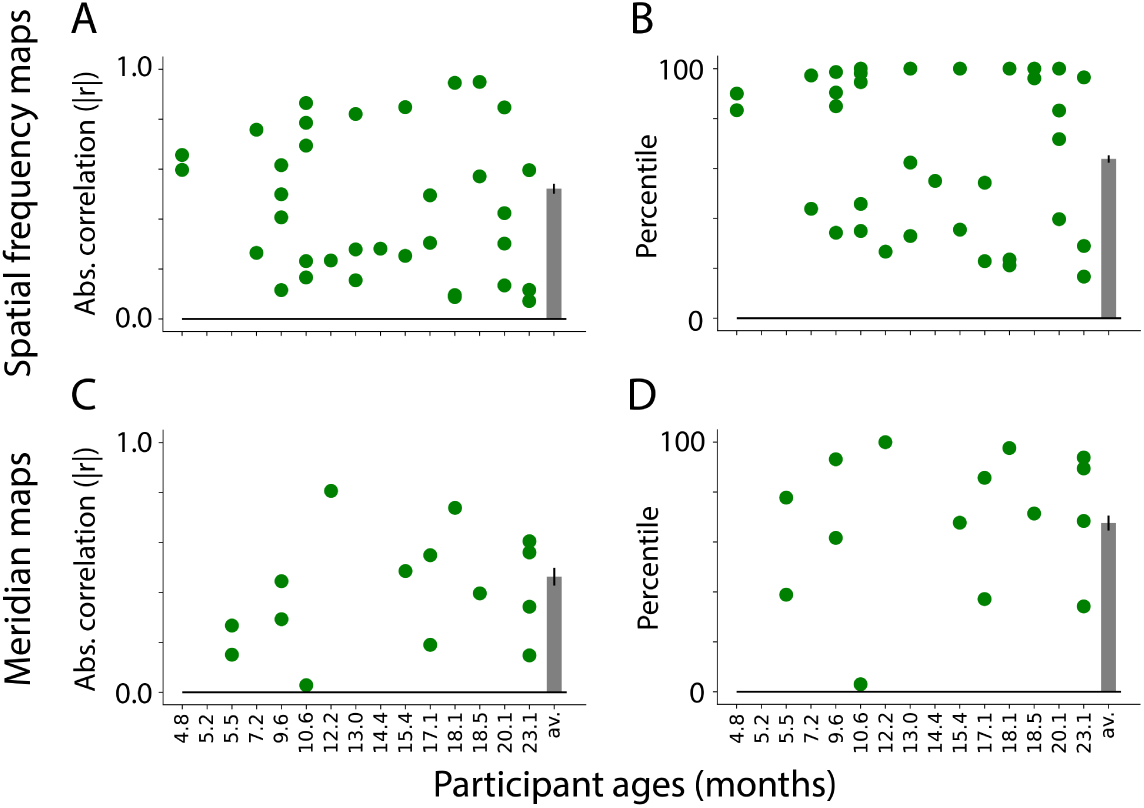
Similarity between visual maps from the retinotopy task and ICA applied to movies. A) Absolute correlation between the task-evoked and component spatial frequency maps (absolute values used because sign of ICA maps is arbitrary). Each dot is a manually identified component. At least one component was identified in 13 out of 15 participants. The bar plot is the average across participants. The error bar is the standard error across participants. B) Ranked correlations for the manually identified spatial frequency components relative to all components identified by ICA. Bar plot is same as A. C) Same as A but for meridian maps. At least one component was identified in 9 out of 15 participants. D) Same as B but for meridian maps.

We performed the same analyses on the meridian maps. As noted above, the lines were now traced perpendicular to the boundaries. Figure 4C shows the correlation between the task-evoked meridian maps and the manually identified components (M=0.46, range: 0.03–0.81). Compared to all possible components identified by ICA, the best possible component was identified for 1 out of 9 participants (Figure 4D). Although the percentile of the average manually identified component was numerically high (M=67.6 percentile, range: 3.0–100.0), it was not significantly above chance (ΔM=17.6, CI=[-1.8–33.0], p=.074). This difference in performance compared to spatial frequency is also evident in the fact that fewer components were identified as potential meridian maps, and that several participants had no such maps. Even so, some participants have components that are highly similar to the meridian maps (e.g., s8037 1 2 or s6687 1 5 in Figure S8). Because it is possible, albeit less likely, to identify meridian maps from ICA, the structure may be present in the data but more susceptible to noise or gaze variability. Spatial frequency maps have a coarser structure than meridian maps, and are more invariant to fixation, which may explain why they are easier to identify. Equivalent analyses of adult data (Figure S3) support this conclusion: meridian maps are found in fewer adult participants.

Despite the similarity of the identified components to the retinotopic maps, it is possible that the components are noise and this similarity arose by chance. Indeed, given enough patterns of spatially smooth noise, some will resemble retinotopic organization. To test how often components derived from noise are misidentified, we made a version of each component in which the functional data were misaligned with respect to the anatomical data while preserving spatial smoothness. We then intermixed an equal number of these “rolled” components amongst the original components and randomized the order such that a coder would be blind as to whether any given component was rolled or original. The blind coder manually categorized each component as a spatial frequency component, a meridian component, or neither (identical to the steps above). It was not possible to make the coder blind for some participants whose rolled data contained visible clues because of partial voluming. In the 6 participants without such clues, only 1 of the 14 components labeled as spatial frequency or meridian, from 920 total components, was a rolled component. The fact that 13 of 14 selected components (93%) were original was extremely unlikely to have occurred by chance (binomial test: *p*=0.002). Thus, our selection procedure rarely identified components as retinotopic in realistic noise.

### Evidence of within area organization with shared response modeling

Finally, we investigated whether the organization of visual cortex in one infant can be predicted from movie-watching data in *other* participants using functional alignment^27^. For such functional alignment to work, stimulus-driven responses to the movie must be shared across participants. These analyses also benefit from greater amounts of data, so we expanded the sample in two ways (Table S2): First, we added 71 movie-watching datasets from additional infants who saw the same movies but did not have usable retinotopy data (and thus were not included in the analyses above that compared movie and retinotopy data within participant). Second, we used data from adult participants, including 8 participants who completed the retinotopy task and saw a subset of the movies we showed infants, and 41 datasets from adults who had seen the movies shown to infants but did not have retinotopy data.

With this expanded dataset, we used shared response modeling (SRM)^33^ to predict visual maps from other participants (Figure 5). Specifically, we held out one participant for testing purposes and used SRM to learn a low-dimensional, shared feature space from the movie-watching data of the remaining participants in a mask of occipital cortex. This shared space represented the responses to that movie in visual cortex that were shared across participants, agnostic to the precise localization of these responses across voxels in each individual (Figure 5A). The number of features in the shared space (K=10) was determined via a cross-validation procedure on movie-watching data in adults (Figure S9). The task-evoked retinotopic maps from all but the held-out participant were transformed into this shared space and averaged, separately for each map type (Figure 5B). We then mapped the held-out participant’s movie data into the learned shared space without changing the shared space (Figure 5C). In other words, the shared response model was learned and frozen before the held-out participant’s data was considered. This approach has been used and validated in prior SRM studies^44^. Taking the inverse of the held-out participant’s mapping allowed us to transform the averaged shared space representation of visual maps into the held-out participant’s brain space (Figure 5D).

**Figure 5:**
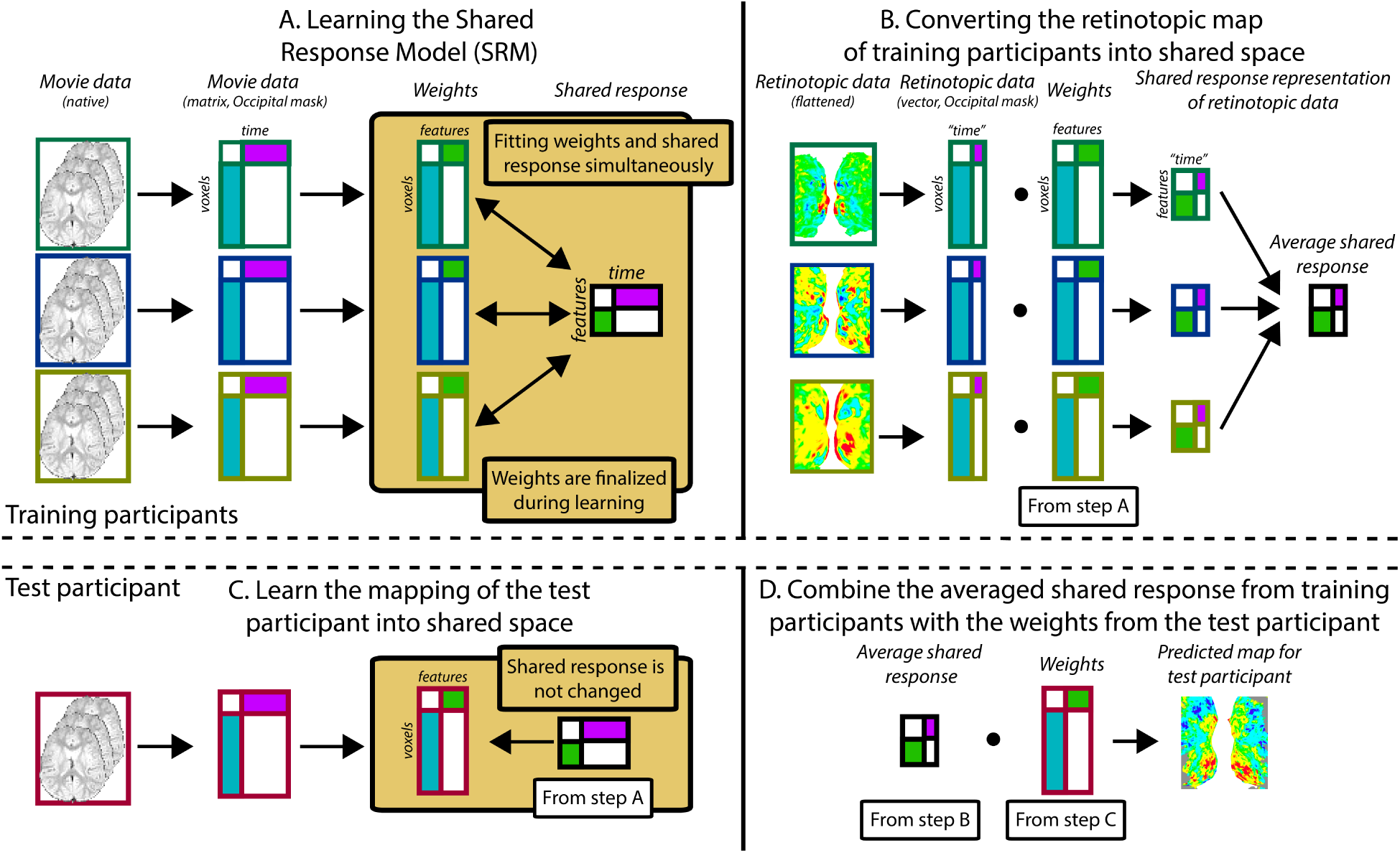
Pipeline for predicting visual maps from movie data. The figure divides the pipeline into 4 steps. All participants watched the same movie. To predict infant data from other infants (or adults), one participant was held out of the training and used as the test participant. Step A: The training participants’ movie data (three color-coded participants shown in this schematic) is masked to include just Occipital voxels. The resulting matrix is run through shared response modeling (SRM)^33^ to find a lower-dimensional embedding (i.e., a weight matrix) of their shared response. Step B: The training participants’ retinotopic maps are transformed into the shared response space using the weight matrices determined in step A. Step C: Once steps A and B are finished, the test participant’s movie data are mapped into the shared space that was fixed from step A. This creates a weight matrix for this test participant. Step D: The averaged shared response of the retinotopic maps from step B is combined with the test participant’s weight matrix from step C to make a prediction of the retinotopic map in the test participant. This prediction can then be validated against their real map from the retinotopy task. Individual gradients for each participant are shown in Figures S10, S11, S12, S13.

This predicted visual organization was compared to the participant’s actual visual map from the retinotopy task using the same methods as for ICA. In other words, the manually traced lines were used to measure the intensity gradients in the predicted maps, and these gradients were com-pared to the ground truth. Critically, predicting the retinotopic maps used no retinotopy data from the held-out participant. Moreover, it is completely unconstrained anatomically (except for a liberal occipital lobe mask). Hence, the similarity of the SRM-predicted map to the task-evoked map is due to representations of visual space in other participants being mapped into the shared space.

We trained SRMs on two populations to predict a held-out infant’s maps: (1) other infants and (2) adults. There may be advantages to either approach: infants are likely more similar to each other than adults in terms of how they respond to the movie; however, their data is more contaminated by motion. When using the infants to predict a held out infant, the spatial frequency map (Figure 6A) and meridian map (Figure 6C) predictions are moderately correlated with task-evoked retinotopy data (spatial frequency: M=0.46, range:-0.06-0.78; meridian: M=0.24, range:-0.12-0.78). Some participants were fit well using SRM (e.g., s2077 1 1, and s6687 1 5 for Figures S10, S11

**Figure 6:**
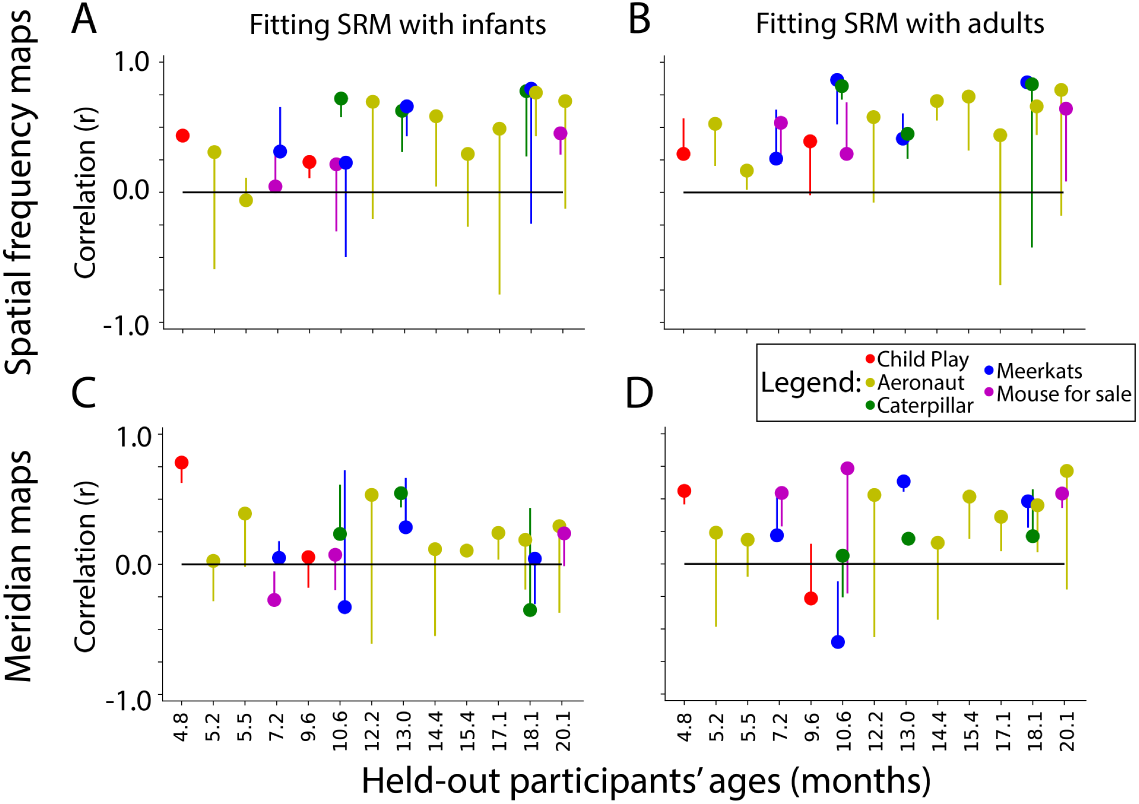
Similarity of SRM-predicted maps and task-evoked retinotopic maps. Correlation be-tween the gradients of the A) spatial frequency maps and C) meridian maps predicted with SRM from other infants and task-evoked retinotopy maps. B, D) Same as A, except using adult partici-pants to train the SRM and predict maps. Dot color indicates the movie used for fitting the SRM. The end of the line indicates the correlation of the task-evoked retinotopy map and the predicted map when using flipped training data for SRM. Hence, lines extending below the dot indicate that the true performance was higher than a baseline fit.

To evaluate whether success was due to fitting the shared response, we flipped the held-out participant’s movie data (i.e., the first timepoint became the last timepoint and vice versa) so that an appropriate fit is not be learnable. The vertical lines for each movie in Figure 6 indicate the change in performance for this baseline. Indeed, flipping significantly worsened prediction of the spatial frequency map (Δ*_Fisher Z_* M=0.52, CI=[0.24–0.80], p*<*.001) and the meridian map (Δ*_Fisher Z_* M=0.24, CI=[0.02–0.49], p=.034). Hence, the movie-evoked response enables the mapping of other infants’ retinotopic maps into a held-out infant.

Using adult data to predict infant data also results in maps similar to task-evoked spatial frequency maps (Figure 6B; M=0.56, range: 0.17–0.79) and meridian maps (Figure 6D; M=0.34, range:-0.27-0.64). Some participants were well predicted by these methods (e.g., s8037 1 2, and s6687 1 4 for Figures S12, S13). Again, flipping the held-out participants movie data significantly worsened prediction of the held-out participant’s spatial frequency map (Δ*_Fisher Z_*M=0.40, CI=[0.17–0.65], p*<*.001) and meridian map (Δ*_Fisher Z_*M=0.33, CI=[0.12–0.55], p=.002). There was no significant difference in SRM performance when using adults versus infants as the training set (spatial frequency: Δ*_Fisher Z_*M=0.14, CI=[-0.00–0.27], p=.054; meridian: Δ*_Fisher Z_* M=0.11, CI=[-0.05–0.28], p=.179). In sum, SRM could be used to predict visual maps with moderate accuracy. This indicates that functional alignment methods like SRM can partially capture the retinotopic organization of visual cortex from infant movie-watching data.

We performed an anatomical alignment analog of the functional alignment (SRM) approach.

This analysis serves as a benchmark for predicting visual maps using task-based data, rather than movie data, from other participants. For each infant participant, we aggregated all other infant or adult participants as a reference. The retinotopic maps from these reference participants were anatomically aligned to the standard surface template, and then averaged. These averages served as predictions of the maps in the test participant, akin to SRM, and were analyzed equivalently (i.e., correlating the gradients in the predicted map with the gradients in the task-based map). These correlations (Table S4) are significantly higher than for functional alignment (using infants to predict spatial frequency, anatomical alignment *<* functional alignment: Δ*_Fisher Z_* M=0.44, CI=[0.32–0.58], p*<*.001; using infants to predict meridians, anatomical alignment < functional alignment:

Δ*_Fisher Z_*M=0.61, CI=[0.47–0.74], p*<*.001; using adults to predict spatial frequency, anatomical alignment < functional alignment: Δ*_Fisher Z_* M=0.31, CI=[0.21–0.42], p*<*.001; using adults to predict meridians, anatomical alignment < functional alignment: Δ*_Fisher Z_* M=0.49, CI=[0.39–0.60], p*<*.001). This suggests that even if SRM shows that movies *can* be used to produce retinotopic maps that are significantly similar to a participant, these maps are not as good as those that can be produced by anatomical alignment of the maps from other participants without any movie data.

## Discussion

We present evidence that movies can reveal the organization of infant visual cortex at different spatial scales. We found that movies evoke differential function across areas, topographic organization of function within areas, and this topographic organization is shared across participants.

We show that the movie-evoked response in a visual area is more similar to the same area in the other hemisphere than to different areas in the other hemisphere. This suggests that visual areas are functionally differentiated in infancy and that this function is shared across hemispheres^31^. By comparing across anatomically distant hemispheres, we reduced the impact of spatial auto-correlation and isolated the stimulus-driven signals in the brain activity^31, 35, 45^. The greater across-hemisphere similarity for same versus different areas provides some of the first evidence that visual areas and streams are functionally differentiated in infants as young as 5 months old. Previous work suggests that functions of the dorsal and ventral streams are detectable in young infants^46^ but that the localization of these functions is immature^47^. Despite this, we find that the areas of infant visual cortex that will mature into the dorsal and ventral streams have distinct activity profiles during movie watching.

Not only do movies evoke differentiated activity in the infant visual cortex between areas, but movies also evoke fine-grained information about the organization of maps within areas. We used a data-driven approach (ICA) to discover maps that are similar to retinotopic maps in the infant visual cortex. We observed components that were highly similar to a spatial frequency map obtained from the same infant in a retinotopy task. This was also true for the meridian maps, to a lesser degree. This means that the retinotopic organization of the infant brain accounts for a detectable amount of variance in visual activity, otherwise components resembling these maps would not be discoverable. Importantly, the components could be identified without knowledge of these ground-truth maps; however, their moderate similarity to the task-defined maps makes them a poor replacement. One caveat for interpreting these results is that although some of the components are *similar* to a spatial frequency map or meridian map, they could reflect a different kind of visual map. For instance, the spatial frequency map is highly correlated with the eccentricity map^22, 48–50^ (which itself is related to receptive field size). This means it is inappropriate to make strong claims about the underlying function of the components based on their similarity to visual maps alone. Another limitation is that ICA does not provide a scale to the variation: although we find a correlation between gradients of spatial frequency in the ground truth and the selected component, we cannot use the component alone to infer the spatial frequency selectivity of any part of cortex. In other words, we cannot infer units of spatial frequency sensitivity from the components alone. Nonetheless, these results do show that it is possible to discover approximations of visual maps in infants and toddlers with movie-watching data and ICA.

We also asked whether functional alignment^29, 33^ could be used to detect visual maps in infants. Using a shared response model^33^ trained on movie-watching data of infants or adults, we transformed the visual maps of other individuals into a held-out infant’s brain to evaluate the fit to visual maps from a retinotopy task^27^. Like ICA, this was more successful for the spatial frequency maps, but it was still possible in some cases with the meridian maps. This is remarkable because the complex pattern of brain activity underlying these visual maps could be ‘compressed’ by SRM into only 10 dimensions in the shared space (i.e., the visual maps were summarized by a vector of 10 values). The weight matrix that ‘decompressed’ visual maps from this low-dimensional space into the held-out infant was learned from their movie-watching data alone. Hence, success with this approach means that visual maps are engaged during infant movie-watching. Furthermore, this result shows that functional alignment is practical for studies in awake infants that produce small amounts of data^51^. This is initial evidence that functional alignment may be useful for enhancing signal quality, like it has in adults^27, 32, 33^, or revealing changing function over development^44^, which may prove especially useful for infant fMRI^51^. In sum, movies evoke sufficiently reliable activity across infants and adults to find a shared response, and this shared response contains information about the organization of infant visual cortex.

To be clear, we are not suggesting that movies work well enough to *replace* a retinotopy task when accurate maps are needed. For instance, even though ICA found components that were highly correlated with the spatial frequency map, we also selected some components that turned out to have lower correlations. Without knowing the ground truth from a retinotopy task, there would be no way to weed these out. Additionally, anatomical alignment (i.e., averaging the maps from other participants and anatomically aligning them to a held-out participant) resulted in maps that were highly similar to the ground truth. Indeed, we previously^23^ found that adult-defined visual areas were moderately similar to infants. While functional alignment with adults can outperform anatomical alignment methods in similar analyses^27^, here we find that functional alignment is inferior to anatomical alignment. Thus, if the goal is to define visual areas in an infant that lacks task-based retinotopy, anatomical alignment of other participants’ retinotopic maps is superior to using movie-based analyses, at least as we tested it.

In conclusion, movies evoke activity in infants and toddlers that recapitulate the organization of the visual cortex. This activity is differentiated across visual areas and contains information about the visual maps at the foundation of visual processing. The work presented here is another demonstration of the power of content-rich, dynamic, and naturalistic stimuli to reveal insights in cognitive neuroscience.

## Methods

### Participants

Infant participants with retinotopy data were previously reported in another study^23^. Of those 17 original sessions, 15 had usable movie data collected in the same session and thus could be included in the current study. In this subsample, the age range was 4.8–23.1 months (*M*=13.0; 12 female; Table S1). The combinations of movies that infants saw were inconsistent, so the types of comparisons vary across analyses reported here. In brief, all possible infant participant sessions (15) were used in the Homotopy analyses and ICA, whereas two of these sessions (ages = 18.5, 23.1 months) could not be used in the SRM analyses. Table S1 reports demographic information for the infant participants. Table S2 reports participant information about each of the movies. It also reports the number and age of participants that were used to bolster the SRM analyses.

An adult sample was collected (N=8, 3 females) and used for validating the analyses and for supporting SRM analyses in infants. Each participant had both retinotopy and movie watching data. The adult participants saw the five most common movies that were seen by infants in our retinotopy sample. To support the SRM analyses, we also utilized any other available adult data from sessions in which we had shown the main movies in otherwise identical circumstances (Table S2).

Participants were recruited through fliers, word of mouth, or the Yale Baby School. This study was approved by the Human Subjects Committee at Yale University. Adults provided informed consent for themselves or their child.

### Materials

Our experiment display code can be found here: https://github.com/ntblab/experiment menu/tree/Movies/ and https://github.com/ntblab/experiment menu/tree/retinotopy/. The code used to perform the data analyses is available at https://github.com/ntblab/infant neuropipe/tree/predict retinotopy/; this code uses tools from the Brain Imaging Analysis Kit^52^; https://brainiak.org/docs/). Raw and preprocessed functional and anatomical data is available here: https://doi.org/10.5061/ dryad.jm63xsjm3.

### Data acquisition

Data were collected at the Brain Imaging Center (BIC) in the Faculty of Arts and Sciences at Yale University. We used a Siemens Prisma (3T) MRI and only the bottom half of the 20-channel head coil. Functional images were acquired with a whole-brain T2* gradient-echo EPI sequence (TR=2s, TE=30ms, flip angle=71, matrix=64×64, slices=34, resolution=3mm iso, interleaved slice acquisition). Anatomical images were acquired with a T1 PETRA sequence for infants (TR1=3.32ms, TR2=2250ms, TE=0.07ms, flip angle=6, matrix=320×320, slices=320, resolution=0.94mm iso, radial slices=30000) and a T1 MPRAGE sequence for adults, with the top of the head coil attached, (TR=2400ms, TE=2.41ms, TI=1000ms, flip angle=8, iPAT=2, slices=176, matrix=256×256, resolution=1.0mm iso).

### Procedure

Our approach for collecting fMRI data from awake infants has been described in a previous methods paper^6^, with important details repeated below. Infants were first brought in for a mock scanning session to acclimate them and their parent to the scanning environment. Scans were scheduled when the infants were typically calm and happy. Participants were carefully screened for metal. We applied hearing protection in three layers for the infants: silicon inner ear putty, over-ear adhesive covers and ear muffs. For the infants that were played sound (see below), Optoacoustics noise canceling headphones were used instead of the ear muffs. The infant was placed on a vacuum pillow on the bed that comfortably reduced their movement. The top of the head coil was not placed over the infant in order to maintain comfort. Stimuli were projected directly onto the surface of the bore. A video camera (High Resolution camera, MRC systems) recorded the infant’s face during scanning. Adult participants underwent the same procedure with the following exceptions: they did not attend a mock scanning session, hearing protection was only two layers (earplugs and Optoacoustics headphones), and they were not on a vacuum pillow. Some infants participated in additional tasks during their scanning session.

When the infant was focused, experimental stimuli were shown using Psychtoolbox^53^ for MATLAB. The details for the retinotopy task are explained fully elsewhere^23^. In short, we showed two types of blocks. For the meridian mapping blocks, a bow tie cut-out of a colorful, large, flickering checkerboard was presented in either a vertical or horizontal orientation^54^. For the spatial frequency mapping blocks, the stimuli were grayscale Gaussian random fields of high (1.5 cycles per visual degree) or low (0.05 cycles per visual degree) spatial frequency^35^. For all blocks, a smaller (1.5 visual degree) grayscale movie was played at center to encourage fixation. Each block type contained two phases of stimulation. The first phase consisted of one of the conditions (e.g., horizontal or high) for 20s, followed immediately by the second phase with the other condition of the same block type (e.g., vertical or low, respectively) for 20s. At the end of each block there was at least 6s rest before the start of the next block. Infant participants saw up to 12 blocks of this stimulus, resulting in 24 epochs of stimuli. Adults all saw 12 blocks.

Participants saw a broad range of movies in this study (Table S3), some of which have been reported previously^28, 30^. The movie titled ‘Child Play’ comprises the concatenation of four silent videos that range in duration from 64–143s and were shown in the same order (with 6s in-between). They extended 40.8° wide by 25.5° high on the screen. The other movies were stylistically similar, computer-generated animations that each lasted 180s. These movies extended 45.0° wide by 25.5° high. Some of the movies were collected as part of an unpublished experiment in which we either played the full movie or inserted drops every 10s (i.e., the screen went blank while the audio continued). We included the ‘Dropped’ movies in the Homotopy analyses and ICA (average number of ‘Dropped’ movies per participant: 0.9, range: 0–3); however, we did not include them in the SRM analyses. Moreover, we only included 4 (out of 17) of these movies in the SRM analyses because there were insufficient numbers of infant participants to enable the training of the SRM.

### Gaze Coding

The infant gaze coding procedure for the retinotopy data was the same as reported previously^23^. The gaze coding for the movies was also the same as reported previously^28, 30^. Par-ticipants looked at the screen for an average of 93.7% of the time (range: 78–99) for the movies used in the homotopy and ICA analyses, and 94.5% of the time (range: 82–99) for the movies used in the SRM analyses (Table S1) Adult participants were not gaze coded, but they were monitored online for inattentiveness. One adult participant was drowsy so they were manually coded. This resulted in the removal of four out of the 24 epochs of retinotopy.

### Preprocessing

We used FSL’s FEAT analyses with modifications in order to implement infant-specific preprocessing of the data^6^. If infants participated in other experiments during the same functional run (14 sessions), the data was split to create a pseudorun. Three burn-in volumes were discarded from the beginning of each run/pseudorun when available. To determine the reference volume for alignment and motion correction, the Euclidean distance between all volumes was calculated and the volume that minimized the distance between all points was chosen as reference (the ‘centroid volume’). Adjacent timepoints with greater than 3mm of movement were interpolated. To create the brain mask we calculated the SFNR^55^ for each voxel in the centroid volume. This produced a bimodal distribution reflecting the signal properties of brain and non-brain voxels. We thresholded the brain voxels at the trough between these two peaks. We performed Gaussian smoothing (FWHM=5mm). Motion correction with 6 degrees of freedom was performed using the centroid volume. AFNI’s despiking algorithm attenuated voxels with aberrant timepoints. The data for each movie were *z*-scored in time.

We registered the centroid volume to a homogenized and skull-stripped anatomical volume from each participant. Initial alignment was performed using FLIRT with a normalized mutual information cost function. This automatic registration was manually inspected and then corrected if necessary using mrAlign from mrTools^56^.

The final step common across analyses created a transformation into surface space. Surfaces were reconstructed from iBEAT v2.0^57^. These surfaces were then aligned into standard Buckner40 standard surface space^58^ using FreeSurfer^58^.

Additional preprocessing steps were taken for the SRM analyses. For each individual movie (including each movie that makes up ‘Child Play’), the fMRI data was time-shifted by 4s and the break after the movie finished was cropped. This was done to account for hemodynamic lag, so that the first TR and last TR of the data approximately^59^ corresponded to the brain’s response to the first and last 2s of the movie, respectively.

Occipital masks were aligned to the participant’s native space for the SRM analyses. To produce these, a mapping from native functional space to standard space was determined. This was enabled using non-linear alignment of the anatomical image to standard space using ANTs^60^. For infants, an initial linear alignment with 12 DOF was used to align anatomical data to the age-specific infant template^61^, followed by non-linear warping using diffeomorphic symmetric normalization. Then, we used a predefined transformation (12 DOF) to linearly align between the infant template and adult standard. For adults, we used the same alignment procedure, except participants were directly aligned to adult standard. We used the occipital mask from the MNI structural atlas^62^ in standard space – defined liberally to include any voxel with an above zero probability of being labelled as the occipital lobe – and used the inverted transform to put it into native functional space.

## Analysis

### Retinotopy

For our measure of task-evoked retinotopy in infants, we used the outputs of the retinotopy analyses from our previous paper^23^ that are publicly released. In brief, we performed separate univariate contrasts between conditions in the study (horizontal*>*vertical, high spatial frequency*>*low spatial frequency). We then mapped these contrasts into surface space. Then, in surface space rendered by AFNI^63^, we demarcated the visual areas V1, V2, V3, V4, and V3A/B using traditional protocols based on the meridian map contrast^64^. We traced lines perpendicular and parallel to the area boundaries to quantify gradients in the visual areas. The anatomically-defined areas of interest^37^ used in Figure 2 were available in this standard surface space. The adult data were also traced using the same methods as infants (described previously^23^) by one of the original infant coders (CE).

### Homotopy

The homotopy analyses compared the time course of functional activity across visual areas in different hemispheres of each infant. For the participants that had more than one movie in a session (N=9), all the movies were concatenated along with burn out time between the movies (Mean number of movies per participant=2.7, range: 1–6, Mean duration of movies=540.7s, range: 186–1116). For the areas that were defined with the retinotopy task (average number of areas traced in each hemisphere = 7.3, range: 6.0–8.0), the functional activity was averaged within area and then Pearson correlated between all other areas. The resulting cross-correlation matrix was Fisher Z transformed before different cells were averaged or compared. If infants did not have an area traced then those areas were ignored in the analyses. We grouped visual areas according to stream, where areas that are *more* dorsal of V1 were called ‘dorsal’ stream and areas *more* ventral were called ‘ventral’ stream. To assess the functional similarity of visual areas, Fisher Z correlations between the same areas in the same stream were averaged, and compared to the correlations of approximately equivalent areas from different streams (e.g., dorsal V2 compared with ventral V2). The averages for each of the two conditions (same stream vs. different stream) were evaluated statistically using bootstrap resampling^65^. Specifically, we computed the mean difference between conditions in a pseudosample, generated by sampling participants with replacement. We created 10,000 such pseudosamples and took the proportion of differences that showed a different sign than the true mean, multiplied by two to get the two-tailed *p*-value. To evaluate how distance affects similarity, we additionally compared with bootstrap resampling the Fisher Z correlations of areas across hemispheres in the same stream: same area to adjacent areas (e.g., ventral V1 with ventral V2), to distal areas (e.g., ventral V1 with ventral V3). Before reporting the results in the figures, the Fisher Z values were converted back into Pearson correlation values.

As an additional analysis to the one described above, we used an atlas of anatomically-defined visual areas from adults^37^ to define both early and later visual areas. Specifically, we used the areas labeled as part of the ventral and dorsal stream (excluding the intraparietal sulcus and frontal eye fields since they often cluster separately^38^), and then averaged the functional response within each area. The functional responses were then correlated across hemispheres, as in the main analysis. Multi-dimensional scaling was then performed on the cross-correlation matrix, and the dimensionality that fell below the threshold for stress (0.2) was chosen. In this case, that was a dimensionality of 2 (stress=0.076). We then visualized the resulting output of the data in these two dimensions.

### Independent Component Analysis (ICA)

To conduct ICA, we provided the preprocessed movie data to FSL’s MELODIC^41^. Like in the homotopy analyses, we used all of the movie data available per session. The algorithm found a range of components across participants (M=76.4 components, range: 31–167). With this large number of possible components, an individual coder (CE) sorted through them to determine whether each one looked like a meridian map, spatial frequency map, or neither (critically, without referring to the ground truth from the retinotopy task). We initially visually inspected each component in volumetric space, looking for the following features: First, we searched for whether there was a strong weighting of the component in visual cortex. Second, we looked for components that had a symmetrical pattern in visual cortex between the two hemispheres. To identify the spatial frequency maps, we looked for a continuous gradient emanating out from the early visual cortex. For meridian maps, we looked for sharp alternations in the sign of the component, particularly near the midline of the two hemispheres. Based on these criteria, we then chose a small set of components that were further scrutinized in surface space. On the surface, we looked for features that clearly define a visual map topography. Again, this selection process was blind to the task-evoked retinotopic maps, so that a person without retinotopy data could take the same steps and potentially find maps. For the adult participants who were analyzed, the components were selected *before* those participants were retinotopically traced, in order to minimize the potential contamination that could occur when performing these manual steps close in time.

These components were then tested against that participant’s task-evoked retinotopic maps. If the component was labeled as a potential spatial frequency map, we tested whether there was a monotonic gradient from fovea to periphery. Specifically, we measured the component response along lines drawn parallel to the area boundaries, averaged across these lines, and then correlated this pattern with the same response in the actual map. The absolute correlation was used because the sign of ICA is arbitrary. For each participant, we then ranked the components to ask if the ones that were chosen were the best ones possible out of all those derived from MELODIC. To test whether the identified components were better than the non-identified components, we ranked all the components correlation to the task-evoked maps. This ranking was converted into a percentile, where 100% means it is the best possible component. We took the identified component’s percentile (or averaged the percentiles if there were multiple components chosen) and compared it to chance (50%). This difference from chance was used for bootstrap resampling to evaluate whether the identified components were significantly better than chance. We performed the same kind of analysis for meridian maps, except in this case the lines used for testing were those drawn perpendicular to the areas. In this case, we were testing whether the components showed oscillations in the sign of the intensity.

To evaluate whether components resembling retinotopic maps arise by chance, we mis-aligned the functional and anatomical data for a subset of participants and manually relabeled them. If retinotopic components are identified at the same rate in the misaligned data as the original data, this would support the concern that the selection process finds structure where there is none. For each participant, we aligned the components to standard surface space and then flipped the labels for left and right hemispheres. Loading these flipped files as if they were correctly aligned had the effect of rolling the functional signals with respect to the anatomy of the cortical surface. Specifically, because the image files are always read in the same order, but the hemispheres differ in the mosaic alignment of nodes in surface space, this flipping transposed voxels from early visual cortex laterally to the approximate position of the lateral occipital cortex and vice versa, while preserving smoothness. Of the 15 total participants, 9 were excluded from this analysis because they had partial volumes (e.g., missing the superior extent of the parietal lobe) such that their rolled data in surface space contained tell-tale signs (e.g., missing voxels were now in an unrealistic place) that precluded blind coding.

To set up the blind test for the coder, the rolled components from the 6 remaining participants were intermixed with an equal number of their original components and then the labels (as original or rolled) were hashed. A coder was given all of the original and rolled components with their hashed names and categorized each one as a spatial frequency component, meridian component, or neither. Once completed, these responses were cross-referenced against the unhashed names to determine whether the components the coder selected as retinotopic had been rolled. The proportion of selected components that were original (vs. rolled) was compared against chance (50%) with a binomial test.

### Shared Response Modeling (SRM)

We based our SRM analyses on previous approaches using hyperalignment^27^ and adapted them for our sample. SRM embeds the brain activity of multiple individuals viewing a common stimulus into a shared space with a small number of feature dimensions. Each voxel of each participant is assigned a weight for each feature. The weight reflects how much the voxel loads onto that feature. For our study, the SRM was either trained on infant movie-watching data or adult movie-watching data to learn the shared response, and the mapping of the training participants into this shared space. For the infant SRM, we used a leave-one-out approach. We took a movie that the held-out infant saw (e.g., ‘Aeronaut’) and considered all other infant participants that saw that movie (including additional participants without any retinotopy data). We fit an SRM model on all of the participants except the held-out one. This model has 10 features, as was determined based on cross-validation with adult data (Figure S9). We used an occipital anatomical mask to fit the SRM. Using the learned individual weight matrices, the retinotopic maps from the infants in the training set were then transformed into the shared space and averaged across participants. The held-out participant’s movie data were used to learn a mapping to the learned SRM features. By applying the inverse of this mapping, we transformed the averaged visual maps of the training set in shared space into the brain space of the held-out participant to predict their visual maps. Using the same methods as described for ICA above, we compared the task-evoked and predicted gradient responses. These analysis steps were also followed for the adult SRM, with the difference being that the group of participants used to create the SRM model and to create the averaged visual maps were adults. As with the infant SRM, additional adult participants without retinotopy data were used for training. Across both types of analysis, the held-out participant was completely ignored when fitting the SRM, and no retinotopy data went into training the SRM.

To test the benefit of SRM, we performed a control analysis in which we scrambled the movie data from the held-out participant before learning their mapping into the shared space. Specifically, we flipped the timecourse of the data so that the first timepoint became the last, and vice versa. By creating a mismatch in the movie sequence across participants, this procedure should result in meaningless weights for the held-out participant and, in turn, the prediction of visual maps using SRM will fail. We compared ‘real’ and ‘flipped’ SRM procedures by computing the difference in fit (transformed into Fisher Z) for each movie, and then averaging that difference within participant across movies. Those differences were then bootstrap resampled to evaluate significance. We also performed bootstrap resampling to compare the ‘real’ SRM accuracy when using infants versus adults for training.

### Anatomical alignment test

We performed a second type of between-participant analysis in addition to SRM. Specifically, we anatomically aligned the retinotopic maps from other participants to make a prediction of the map in a held-out participant. To achieve this, we first aligned all spatial frequency and meridian maps from infant and adult participants with retinotopy into the Buckner40 standard surface space^58^. For each infant participant, we composed a map from the average of the *other* participants. The other participants were either all the other infants or all the adult participants. We then used the lines traced parallel to the area boundaries (for spatial frequency) or perpendicular to the area boundaries (for meridian) to extract gradients of response in the average maps. These gradients were then correlated with the ground-truth gradients (i.e., the alternations in sensitivity in the held-out infant using lines traced from that participant). These correlations were then compared to SRM results within participants using bootstrap resampling. If a participant had multiple movies worth of data, then they were averaged prior to this comparison.

### Contributions

C.T.E. and M.J.A. conceived of the analyses. C.T.E., T.S.Y., & N.B.T-B. collected the data. C.T.E. & T.S.Y. preprocessed the data. C.T.E. performed the analyses. All authors contributed to the drafting of the manuscript.

## Acknowledgements

We are thankful to the families of infants who participated. We also acknowledge the hard work of the Yale Baby School team, including L. Rait, J. Daniels, A. Letrou, and K. Armstrong for recruitment, scheduling, and administration, and L. Skalaban, A. Bracher, D. Choi, and J. Trach for help in infant fMRI data collection. Thank you to J. Wu, J. Fel, and A. Klein for help with gaze coding, and R. Watts for technical support. We are grateful for internal funding from the Department of Psychology and Faculty of Arts and Sciences at Yale University. N.B.T-B. was further supported by the Canadian Institute for Advanced Research and the James S. McDonnell Foundation (https://doi.org/10.37717/2020-1208).

## Supplementary materials

Movies reveal the fine-grained organization of infant visual cortex

Ellis, C. T., Yates, T. S., Arcaro, M. J., & Turk-Browne, N. B.

**Table S1:**
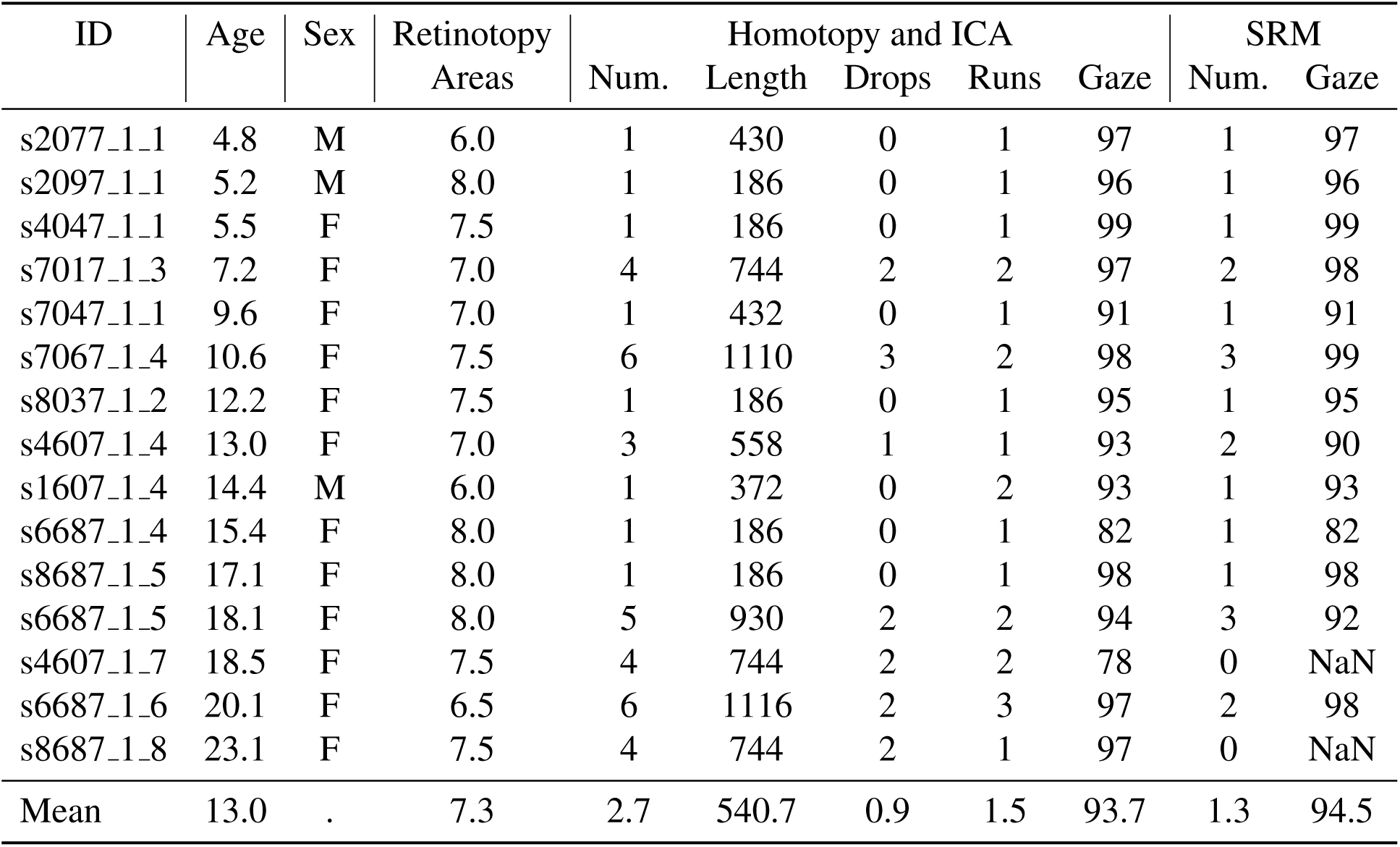
Demographic and dataset information for infant participants in the study. ‘Age’ is recorded in months. ‘Sex’ is the assigned sex at birth. ‘Retinotopy areas’ is the number of areas segmented from task-evoked retinotopy, averaged across hemispheres. Information about the movie data is separated based on analysis type: whereas all movie data is used for homotopy and ICA analyses, a subset of data is used for SRM. ‘Num.’ is the number of movies used. ‘Length’ is the duration in seconds of the run used for these analyses (includes both movie and rest periods). ‘Drops’ is the number of movies that include dropped periods. ‘Runs’ says how many runs or pseudoruns of movie data there were. ‘Gaze’ is the percentage of the data where the participants were looking at the movie.

**Table S2:**
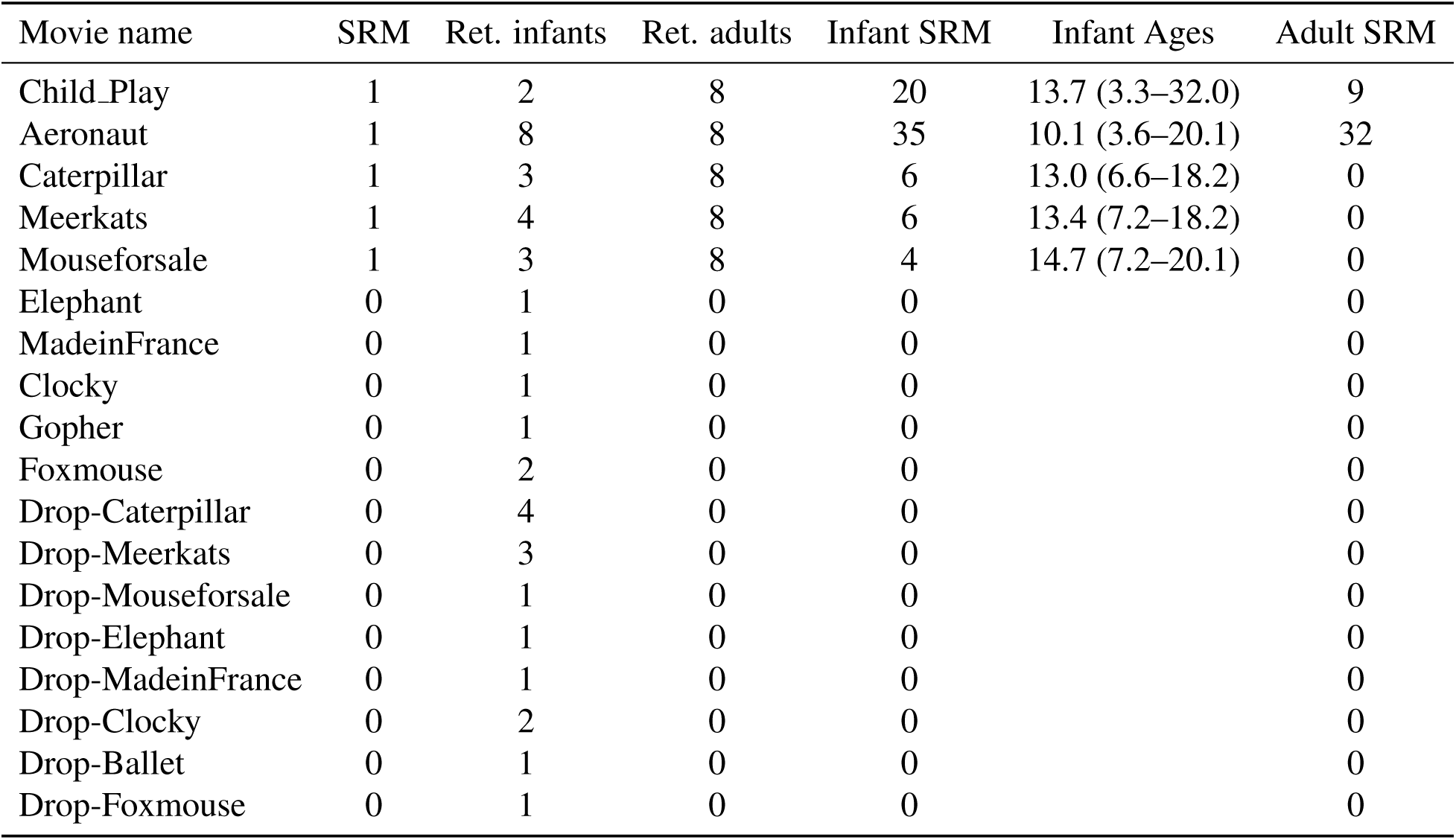
Number of participants per movie. The first column is the movie name, where ‘Drop-’ indicates that it was a movie containing alternating epochs of blank screens. ‘SRM’ indicates whether the movie is used in SRM analyses. The movies that are not included in SRM are used for homotopy and ICA. ‘Ret. infants’ and ‘Ret. adults’ refers to the number of participants with retinotopy data that saw this movie. ‘Infant SRM’ and ‘Adult SRM’ refer to the number of additional participants available to use for training the SRM but who did not have retinotopy data. ‘Infant Ages’ is the average age in months of the infant participants included in the SRM, with the range of ages included in parentheses.

| Movie name | SRM | Ret. infants | Ret. adults | Infant SRM | Infant Ages | Adult SRM |
| --- | --- | --- | --- | --- | --- | --- |
| Child_Play | 1 | 2 | 8 | 20 | 13.7 (3.3–32.0) | 9 |
| Aeronaut | 1 | 8 | 8 | 35 | 10.1 (3.6–20.1) | 32 |
| Caterpillar | 1 | 3 | 8 | 6 | 13.0 (6.6–18.2) | 0 |
| Meerkats | 1 | 4 | 8 | 6 | 13.4 (7.2–18.2) | 0 |
| Mouseforsale | 1 | 3 | 8 | 4 | 14.7 (7.2–20.1) | 0 |
| Elephant | 0 | 1 | 0 | 0 |  | 0 |
| MadeinFrance | 0 | 1 | 0 | 0 |  | 0 |
| Clocky | 0 | 1 | 0 | 0 |  | 0 |
| Gopher | 0 | 1 | 0 | 0 |  | 0 |
| Foxmouse | 0 | 2 | 0 | 0 |  | 0 |
| Drop-Caterpillar | 0 | 4 | 0 | 0 |  | 0 |
| Drop-Meerkats | 0 | 3 | 0 | 0 |  | 0 |
| Drop-Mouseforsale | 0 | 1 | 0 | 0 |  | 0 |
| Drop-Elephant | 0 | 1 | 0 | 0 |  | 0 |
| Drop-MadeinFrance | 0 | 1 | 0 | 0 |  | 0 |
| Drop-Clocky | 0 | 2 | 0 | 0 |  | 0 |
| Drop-Ballet | 0 | 1 | 0 | 0 |  | 0 |
| Drop-Foxmouse | 0 | 1 | 0 | 0 |  | 0 |

**Table S3:** Details for each movie used in this study.‘Name’ specifies the movie name. ‘Duration’ specifies the duration of the movie in seconds. Movies were edited to standardize length and remove inappropriate content. ‘Sound’ is whether sound was played during the movie. These sounds include background music, animal noises, and sound effects, but no language. ‘Description’ gives a brief description of the movie, as well as a current link to it when appropriate. All movies are provided in the data release.

| Name | Duration | Sound | Description |
| --- | --- | --- | --- |
| Child_Play | 406 | 0 | Four photo-realistic clips from ‘Daniel Tiger’ showing children playing. The clips showed the following: 1. children playing in an indoor playground (84s); 2. a family making frozen banana desserts (64s); 3. a child visiting the doctor (115s); 4. children helping with indoor and outdoor chores (143s). |
| Aeronaut | 180 | 0 | A computer-generated segment from a short film titled “Soar” (https://vimeo.com/148198462) and described here<sup>28</sup>. |
| Caterpillar | 180 | 1 | A computer-generated segment from a short film titled “Sweet Cocoon” (https://www.youtube.com/watch?v=yQ1ZcNpbwOA). This video depicts a caterpillar trying to fit into its cocoon so it can become a butterfly. |
| Meerkats | 180 | 1 | A computer-generated segment from the short film titled “Catch It” (https://www.youtube.com/watch?v=c88QE6yGhfM). It depicts a gang of meerkats who take back a treasured fruit from a vulture. |
| Mouse for Sale | 180 | 1 | A computer-generated segment from a short film of the same name (https://www.youtube.com/watch?v=UB3nKCNUBB4). It shows a mouse in a pet store who is teased for having big ears. |
| Elephant | 180 | 1 | A computer-generated segment from a short film of the same name (https://www.youtube.com/watch?v=h_aC8pGY1aY). It shows an elephant in a china shop. |
| Made in France | 180 | 1 | A computer-generated segment from a short film of the same name (https://www.youtube.com/watch?v=Her3d1DH7yU). It shows a mouse making cheese. |
| Clocky | 180 | 1 | A computer-generated segment from a short film of the same name (https://www.youtube.com/watch?v=8VRD5KOFK94). It shows a clock preparing to wake up its owner. |
| Gopher | 180 | 1 | A computer-generated segment named ‘Gopher broke’ (https://www.youtube.com/watch?v=tWufIUbXubY). It shows a gopher collecting food. |
| Foxmouse | 180 | 1 | A computer-generated segment named ‘The short story of a fox and a mouse’ (https://www.youtube.com/watch?v=k6kCwj0Sk4s). It shows a fox playing with a mouse in the snow. |
| Ballet | 180 | 1 | A computer-generated segment named ‘The Duet’ (https://www.youtube.com/watch?v=GuX52wkCIJA). This is an artistic rendition of growing up and falling in love. |

**Table S4:** Correlations between infant gradients and the spatial average of other infants or adults. For each participant, all other participants with retinotopy data (adults or infants) were aligned to standard surface space and averaged. The traced lines from the held-out participant were then applied to this average. The resulting gradients were correlated with the held-out participant and the correlation is reported here. This was done separately for meridian maps and spatial frequency maps.

| ID | Adults |  | Infants |  |
| --- | --- | --- | --- | --- |
|  | Spatial freq. | Meridians | Spatial freq. | Meridians |
| s2077_1_1 | 0.85 | 0.77 | 0.89 | 0.81 |
| s2097_1_1 | 0.66 | 0.72 | 0.66 | 0.65 |
| s4047_1_1 | 0.86 | 0.78 | 0.94 | 0.82 |
| s7017_1_3 | 0.90 | 0.92 | 0.93 | 0.92 |
| s7047_1_1 | 0.43 | 0.65 | 0.56 | 0.64 |
| s7067_1_4 | 0.87 | 0.67 | 0.92 | 0.61 |
| s8037_1_2 | 0.92 | 0.73 | 0.93 | 0.83 |
| s4607_1_4 | 0.77 | 0.97 | 0.74 | 0.94 |
| s1607_1_4 | 0.93 | 0.82 | 0.92 | 0.86 |
| s6687_1_4 | 0.87 | 0.90 | 0.93 | 0.93 |
| s8687_1_5 | 0.97 | 0.89 | 0.98 | 0.83 |
| s6687_1_5 | 0.92 | 0.81 | 0.97 | 0.91 |
| s4607_1_7 | 0.85 | 0.91 | 0.80 | 0.86 |
| s6687_1_6 | 0.92 | 0.94 | 0.86 | 0.97 |
| s8687_1_8 | 0.89 | 0.93 | 0.90 | 0.88 |
| Mean | 0.84 | 0.83 | 0.86 | 0.83 |

---

**Figure S1:**
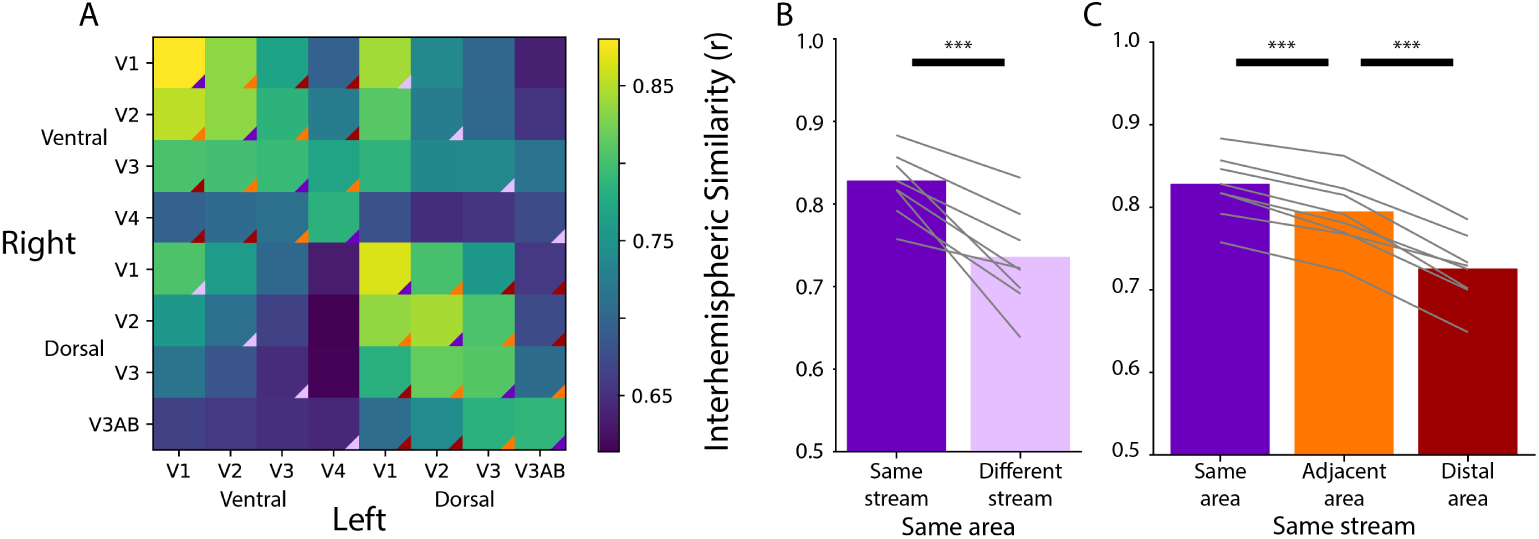
Homotopic correlations between retinotopic areas in the adult sample, akin to Figure 1. (A) Average correlation of the timecourse of activity evoked during movie watching for all areas. Correlation of homotopic areas: M=0.83 (range: 0.78–0.88). (B) Across-hemisphere similarity of the same visual area from the same stream and from different streams. Difference with bootstrap resampling: Δ*_Fisher Z_* M=0.24, p*<*0.001. (C) Across-hemisphere similarity in the same stream when matching the same area, matching to an adjacent area, or matching to a distal area. Difference with bootstrap resampling: Same *>* Adjacent Δ*_Fisher Z_* M=0.10, p*<*0.001; Adjacent *>* Distal Δ*_Fisher Z_* M=0.16, p*<*0.001. Grey lines represent individual participants. *** = p*<*0.001 from bootstrap resampling.

**Figure S2:**
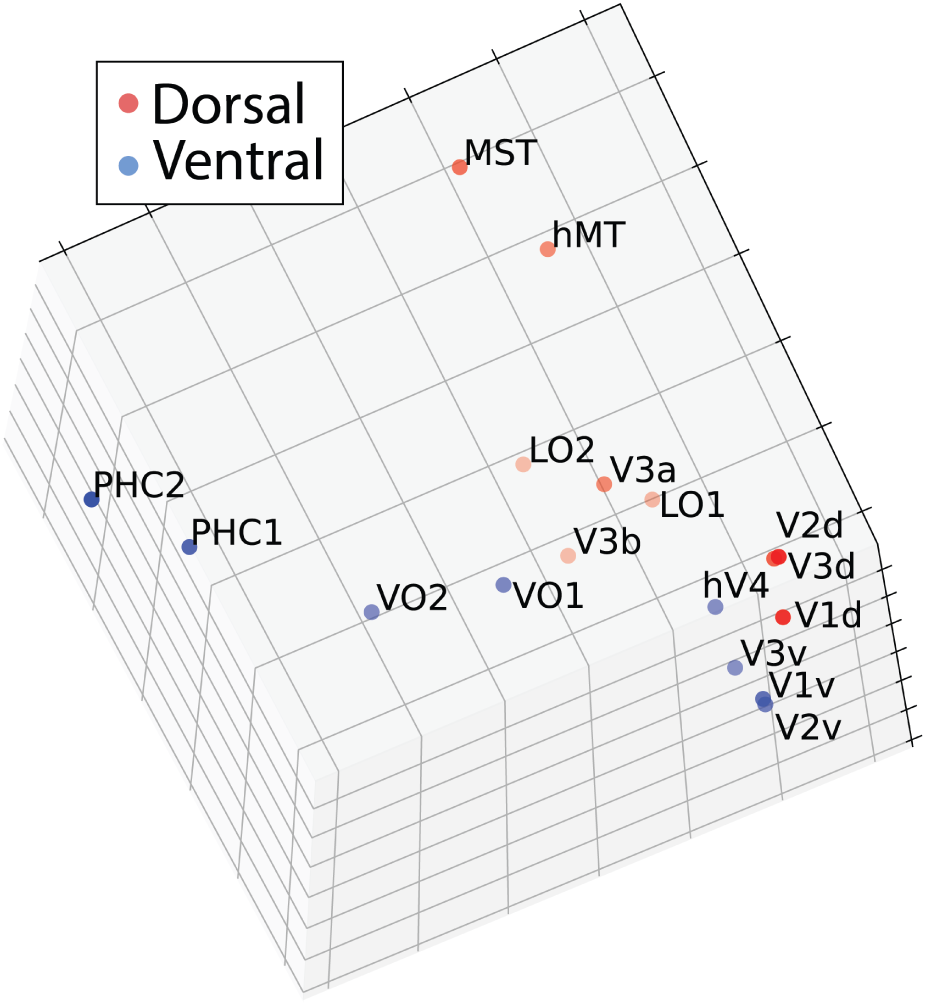
Multi-dimensional scaling of movie-evoked activity in adult visual cortex, akin to Figure 2. A 2-dimensional embedding had inappropriately high stress – 0.87 – whereas a 3-dimensional embedding had appropriate stress: 0.105. This 3-dimensional scatter depicts the similarity of the functional timecourse of areas as a function of Euclidean distance. The plot shows a projection that emphasizes the similarity to the brain’s organization.

**Figure S3:**
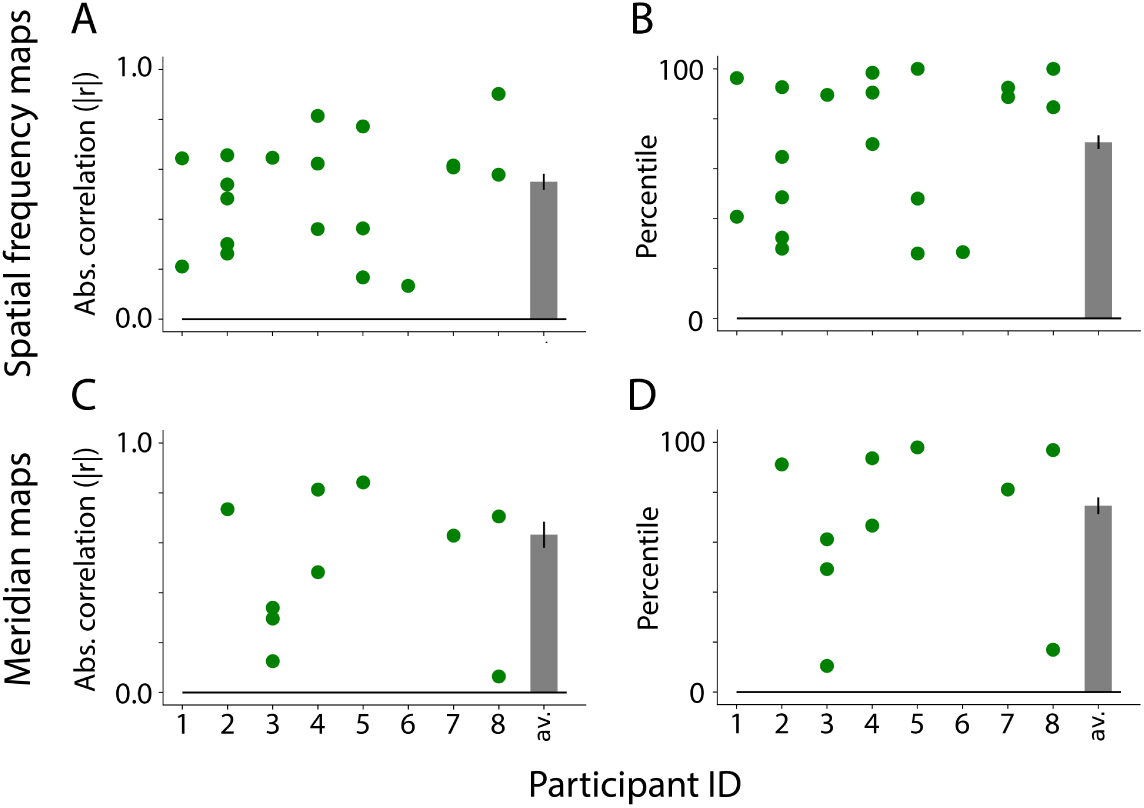
Similarity between visual maps from the adult retinotopy task and ICA applied to movies, akin to Figure 4. A) Absolute correlation between the task-evoked and component spatial frequency maps (absolute values used because sign of ICA maps is arbitrary). Each dot is a manually identified component. At least one component was identified in 8 out of 8 adult participants. The bar plot is the average across participants. The error bar is the standard error across participants. B) Ranked correlations for the manually identified spatial frequency components relative to all components identified by ICA. Bar plot is same as A. Percentile tests: M=70.6 percentile, range: 26.6–92.3, ΔM from chance=20.6, CI=[4.2–34.9], p=.014. C) Same as A but for meridian maps. At least one component was identified in 6 out of 8 participants. D) Same as B but for meridian maps. Percentile tests: M=74.6 percentile, range: 40.3–98.0, ΔM from chance=24.6, CI=[8.2–39.6], p=.004.

**Figure S4:**
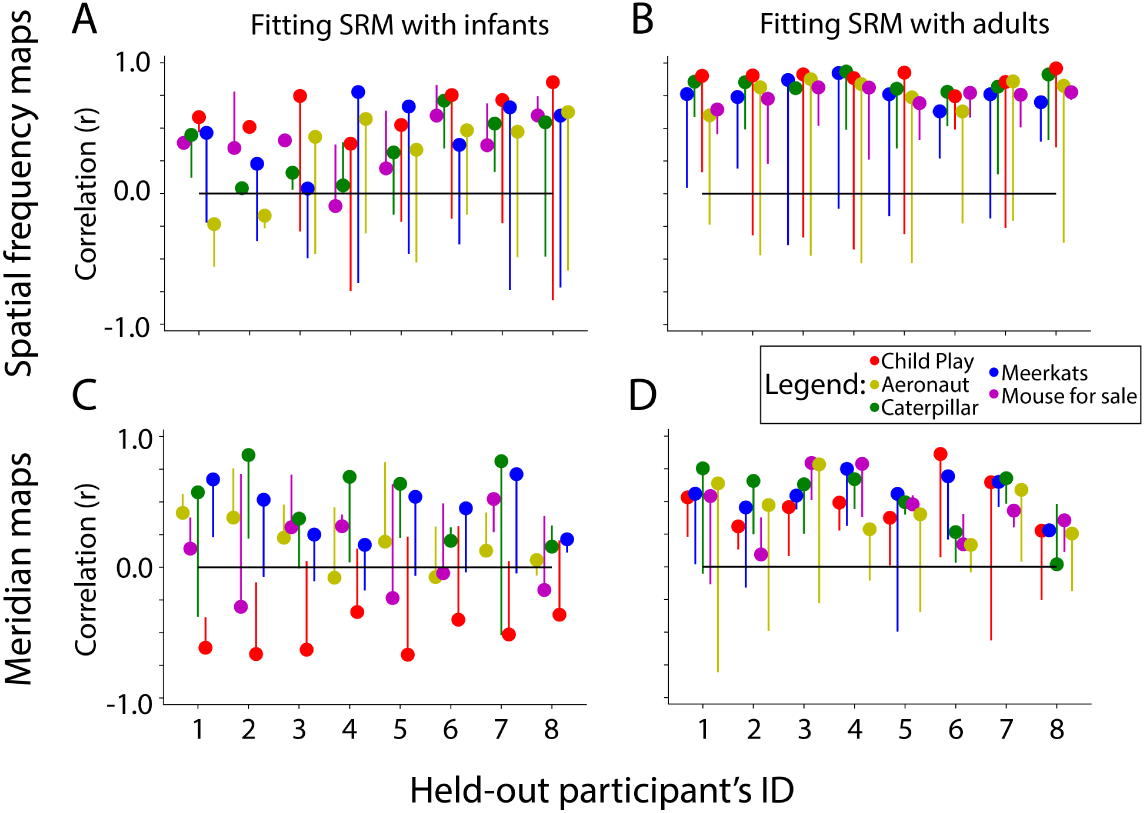
Similarity of SRM-predicted maps and task-evoked retinotopic maps in adults, akin to Figure 6. Correlation between the gradients of the A) spatial frequency maps and C) meridian maps predicted with SRM from infants and their task-evoked retinotopy maps. Difference between real and flipped SRM fit: Spatial frequency= Δ*_Fisher Z_* M=0.59, CI=[0.36–0.83], p*<*.001. Meridian= Δ*_Fisher Z_* M=-0.07, CI=[-0.22–0.10], p=.382. Note: only two infants were used in the prediction with Child Play (red dots), hence why they likely show erratic behavior. B, D) Same as A, except using adult participants to train the SRM and predict maps. Difference between real and flipped SRM fit: Spatial frequency= Δ*_Fisher Z_* M=1.05, CI=[0.85–1.22], p*<*.001. Meridian= Δ*_Fisher Z_* M=0.49, CI=[0.36–0.64], p*<*.001. Dot color indicates the movie used for fitting the SRM. The end of the line indicates the correlation of the task-evoked retinotopy map and the predicted map when using flipped training data for SRM.

**Figure S5:**
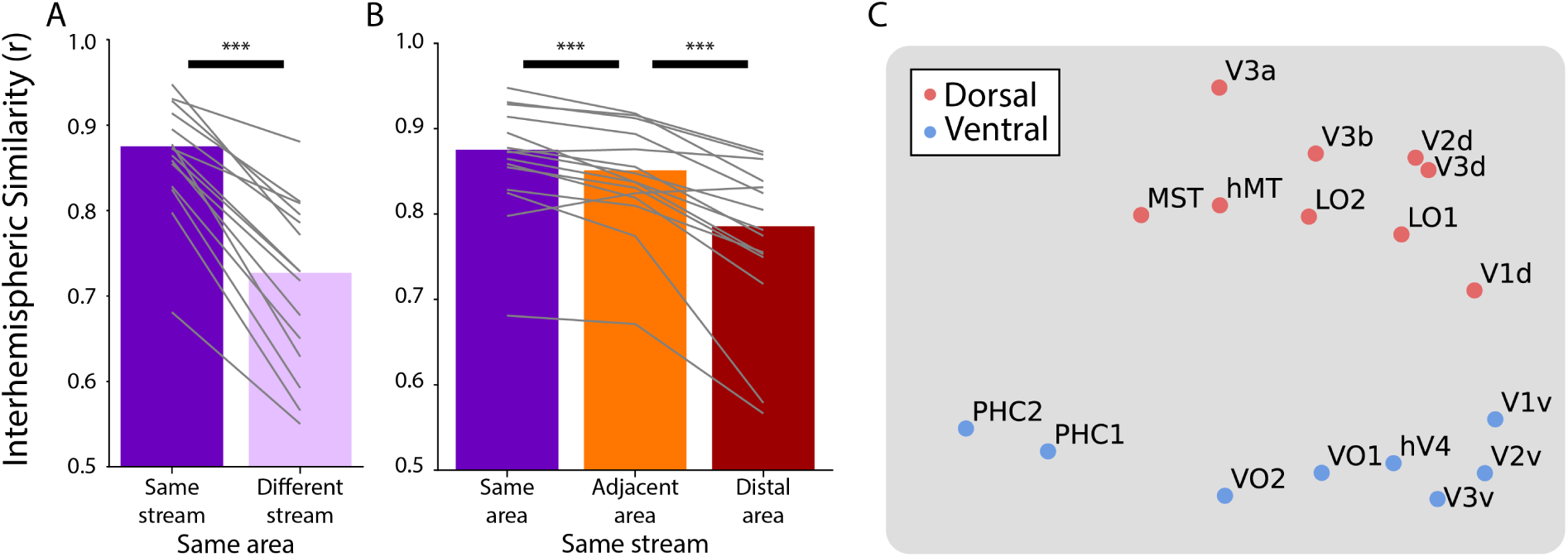
Homotopic correlations when controlling for motion. In this analysis, we computed correlations for all pairwise comparisons while partialing out our metric of motion: framewise displacement. In other words, if the functional timecourse in an area was correlated with the motion metric then this would decrease the correlation between that area and others. Subfigures A and B use task-evoked retinotopic definitions of areas (akin to Figure 1), whereas subfigure C uses anatomical definitions of areas (akin to Figure 2). Overall the results are qualitatively similar, suggesting that motion does not explain the effect observed here. A) Correlation of the same area and same stream (e.g., left ventral V1 and right ventral V1) versus the same area and different stream (e.g., left ventral V1 and right dorsal V1). Difference with bootstrap resampling: Δ*_Fisher Z_* M=0.43, p*<*0.001. B) Correlation within the same stream between the same areas, adjacent areas (e.g., left ventral V1 and right ventral V2), or distal areas (e.g., left ventral V1 and right ventral hV4). Difference with bootstrap resampling: Same *>* Adjacent Δ*_Fisher Z_* M=0.09, p*<*0.001; Adjacent *>* Distal Δ*_Fisher Z_* M=0.20, p*<*0.001. Grey lines represent individual participants. *** = p*<*0.001 from bootstrap resampling. C) Multidimensional scaling of the partial correlation between all anatomically defined areas. The timecourse of functional activity for each area was extracted and correlated across hemispheres, while partialing out framewise displacement. This matrix was averaged across participants and used to create a Euclidean dissimilarity matrix. MDS captured the structure of this matrix in two dimensions with suitably low stress (0.089). The plot shows a projection that emphasizes the similarity to the brain’s organization.

**Figure S6:**
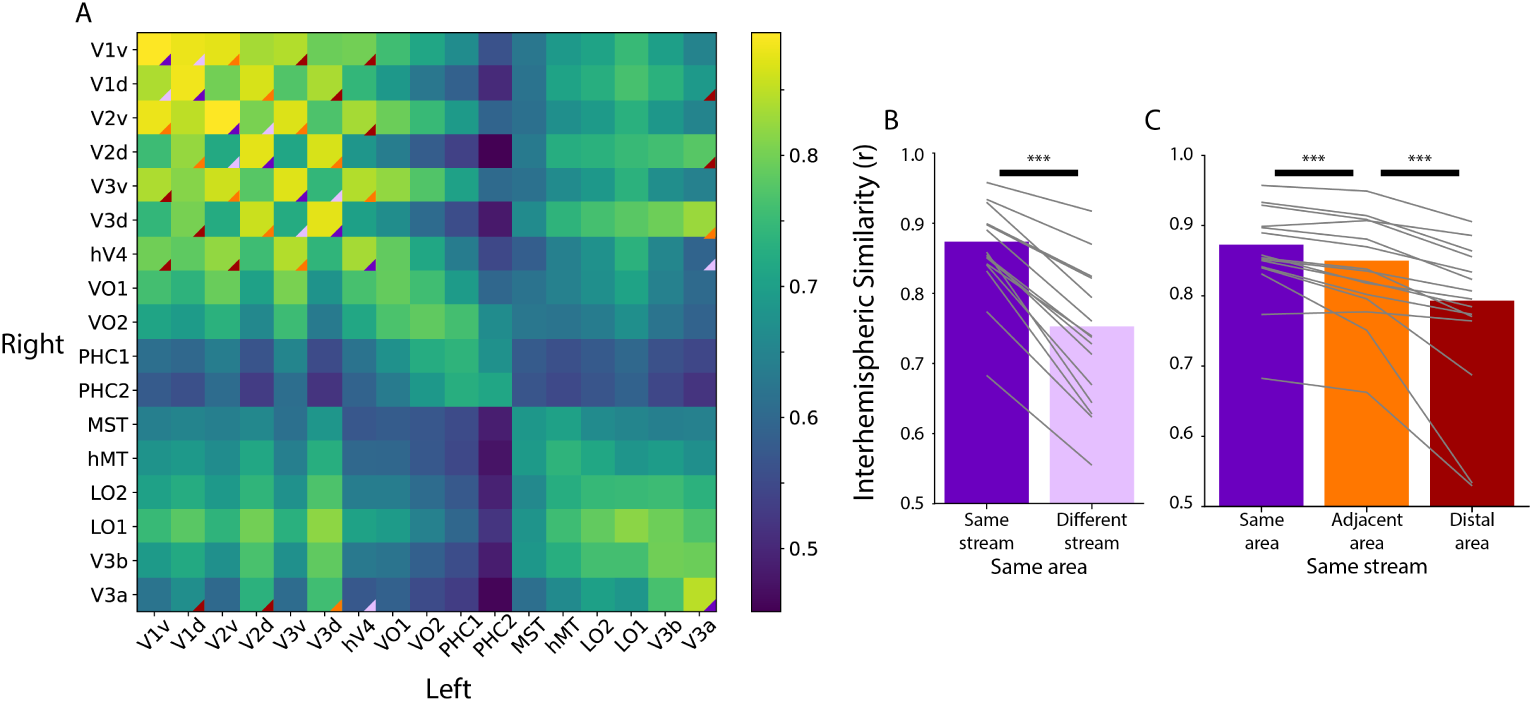
Homotopic correlations between anatomically defined areas corresponding to the data used in Figure 2. A) Average correlation of the time course of activity evoked during movie watching for ventral and dorsal areas in an anatomical segmentation^37^. This is done for the left and right hemispheres separately, which is why the matrix is not diagonally symmetric. The triangles overlaid on the matrix corner highlights the area-wise comparisons used in B and C. Only areas that we were able to retinotopically map (i.e., those that overlap with Figure 1) were used for this analysis. B) Correlation of the same area and same stream (e.g., left ventral V1 and right ventral V1) versus the same area and different stream (e.g., left ventral V1 and right dorsal V1). Difference with bootstrap resampling: Δ*_Fisher Z_* M=0.37, p*<*0.001. C) Correlation within the same stream between the same areas, adjacent areas (e.g., left ventral V1 and right ventral V2), or distal areas (e.g., left ventral V1 and right ventral hV4). Difference with bootstrap resampling: Same *>* Adjacent Δ*_Fisher Z_* M=0.09, p*<*0.001; Adjacent *>* Distal Δ*_Fisher Z_* M=0.18, p*<*0.001. Grey lines represent individual participants. *** = p*<*0.001 from bootstrap resampling

**Figure S7:**
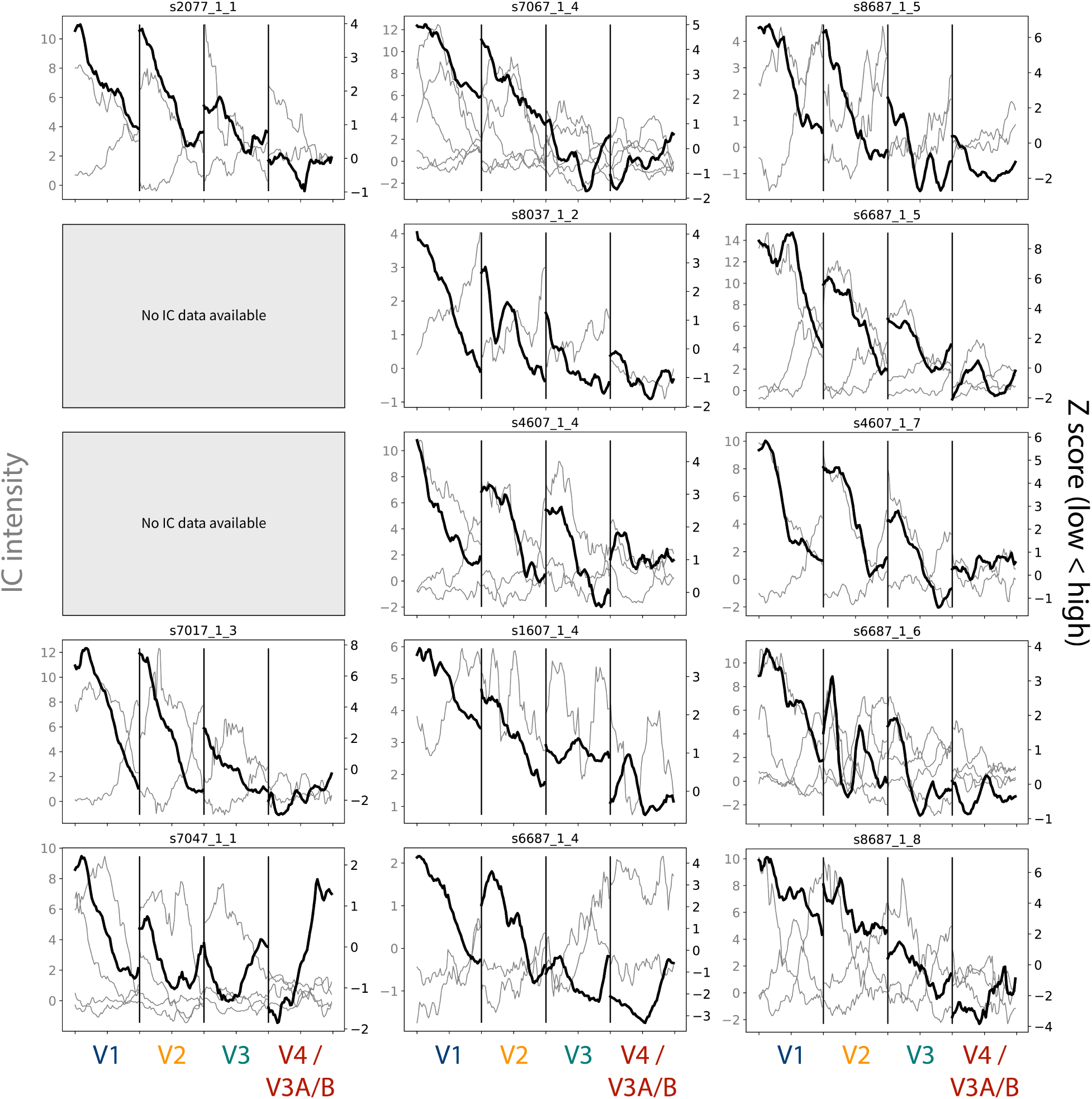
Gradients for the task-evoked and ICA-based spatial frequency maps. The grey lines depict the gradients from each chosen IC map, and their scale is indicated by the Y-axis on the left-hand side. The sign of the maps have not been edited, but it is arbitrary. The black line indicates the gradient from the task-evoked map, and their scale is indicated by the Y-axis on the right-hand side. Participants are listed in order of age. Participant data is not reported if no components were chosen for that participant.

**Figure S8:**
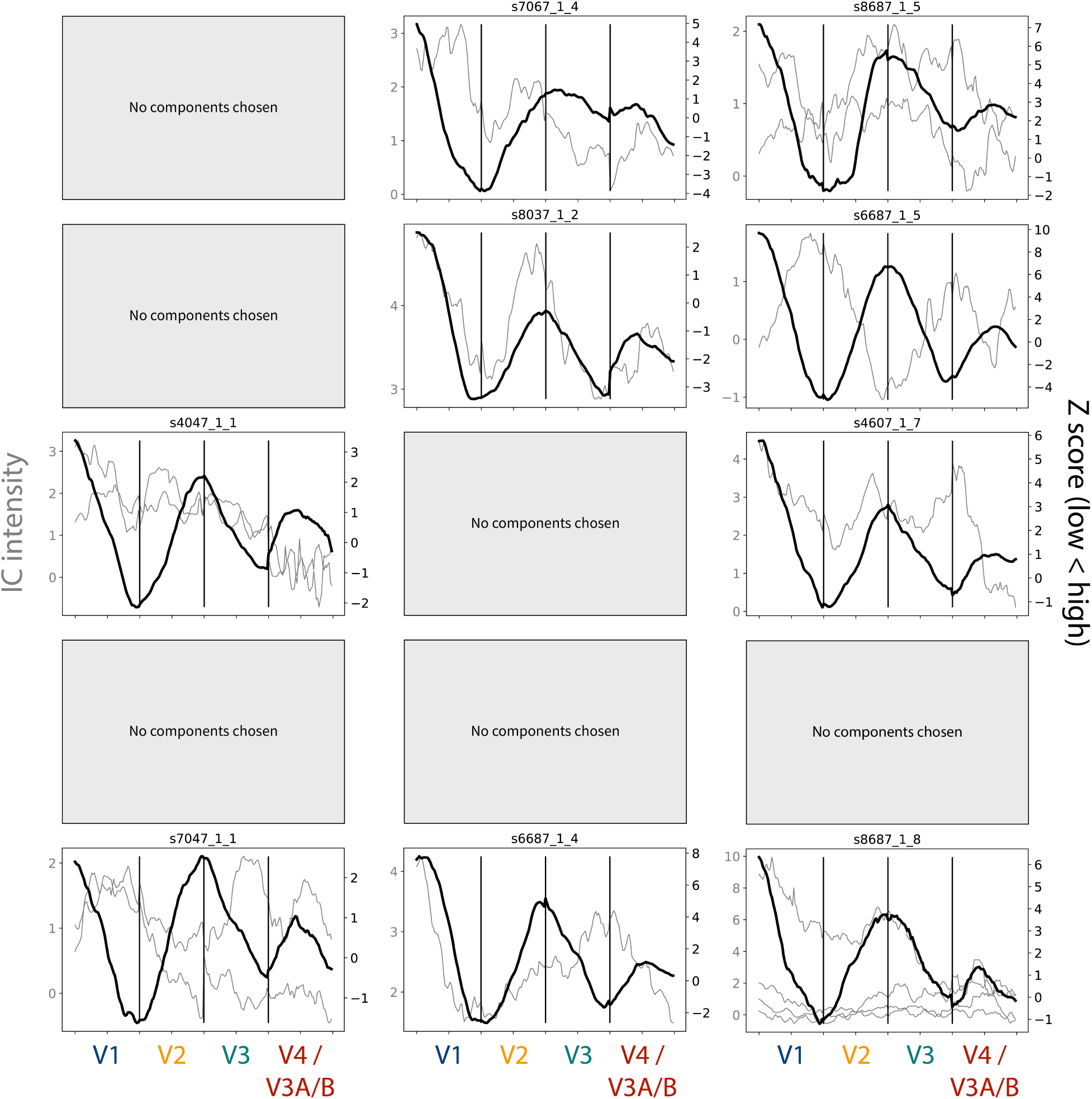
Gradients for the task-evoked and ICA-based meridian maps. The grey lines depict the gradients from each chosen IC map, and their scale is indicated by the Y-axis on the left-hand side. The sign of the maps have not been edited, but it is arbitrary. The black line indicates the gradient from the task-evoked map, and their scale is indicated by the Y-axis on the right-hand side. Participants are listed in order of age. Participant data is not reported if no components were chosen for that participant.

**Figure S9:**
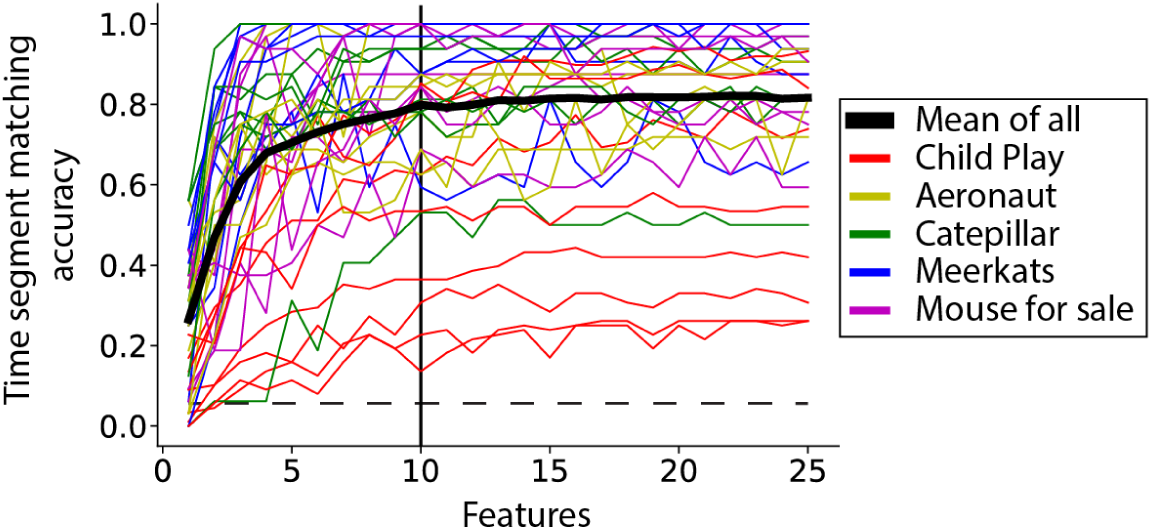
Cross-validation of the number of features in SRM. The movie data from all adult participants (Table S2) was split in half, with a 10 TR buffer between sets. The data were masked only to include occipital lobe voxels. The first half of the movie was used for training the SRM in all but one participant. The number of features learned by the SRM was varied across analyses from 1–25. The second half of the movie was then used to generate a shared response (i.e., the activity time course in each feature). To test the SRM, the held-out participant’s first half of data is used to learn a mapping of that participant into the SRM space (this mapping does not change the features learned and is not based on the second half of data). The second half of the held-out participant’s data is then mapped into the shared response space, like the other participants. Time-segment matching was performed on the shared response^29, 33^. In brief, time-segment matching tests whether a segment of the data (10 TRs) in the held-out participant can be matched to its correct time point based on the other participants. This tests whether the SRM succeeds in making the held-out participant similar to the others. This analysis was performed on each participant and movie separately (each has a line). The dashed line is chance for time-segment matching, averaged across all movies and participants. The black solid line at features=10 reflects the number of features chosen.

**Figure S10:**
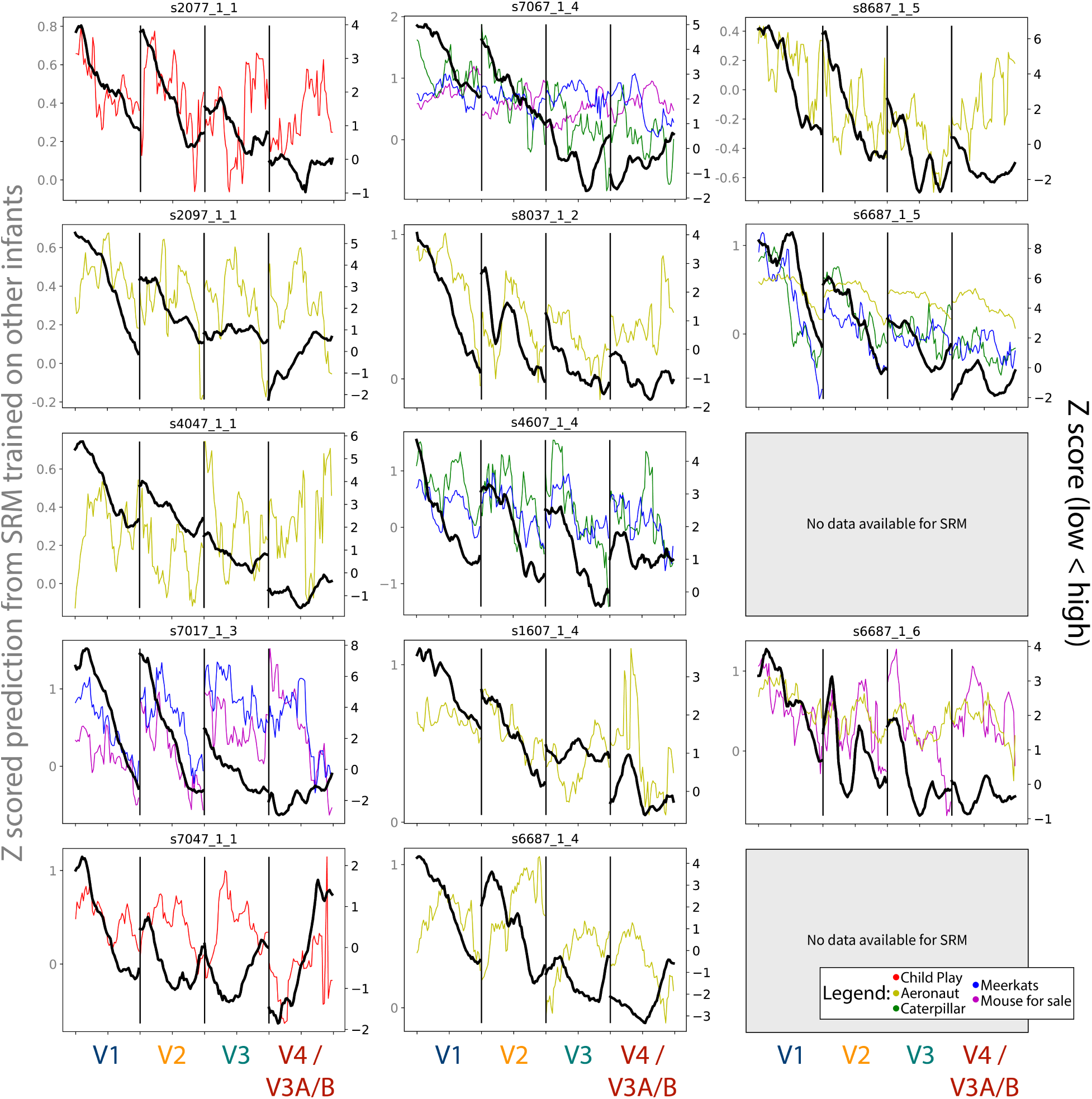
Gradients for the spatial frequency maps predicted using SRM from other infant participants, compared to the task-evoked gradients. The colored lines depict the gradients from each chosen movie that could be used, and their scale is indicated by the Y-axis on the left-hand side. The black line indicates the gradient from the task-evoked map, and their scale is indicated by the Y-axis on the right-hand side. Participants are listed in order of age. Participant data is not reported if the participant did not have SRM-compatible movie data.

**Figure S11:**
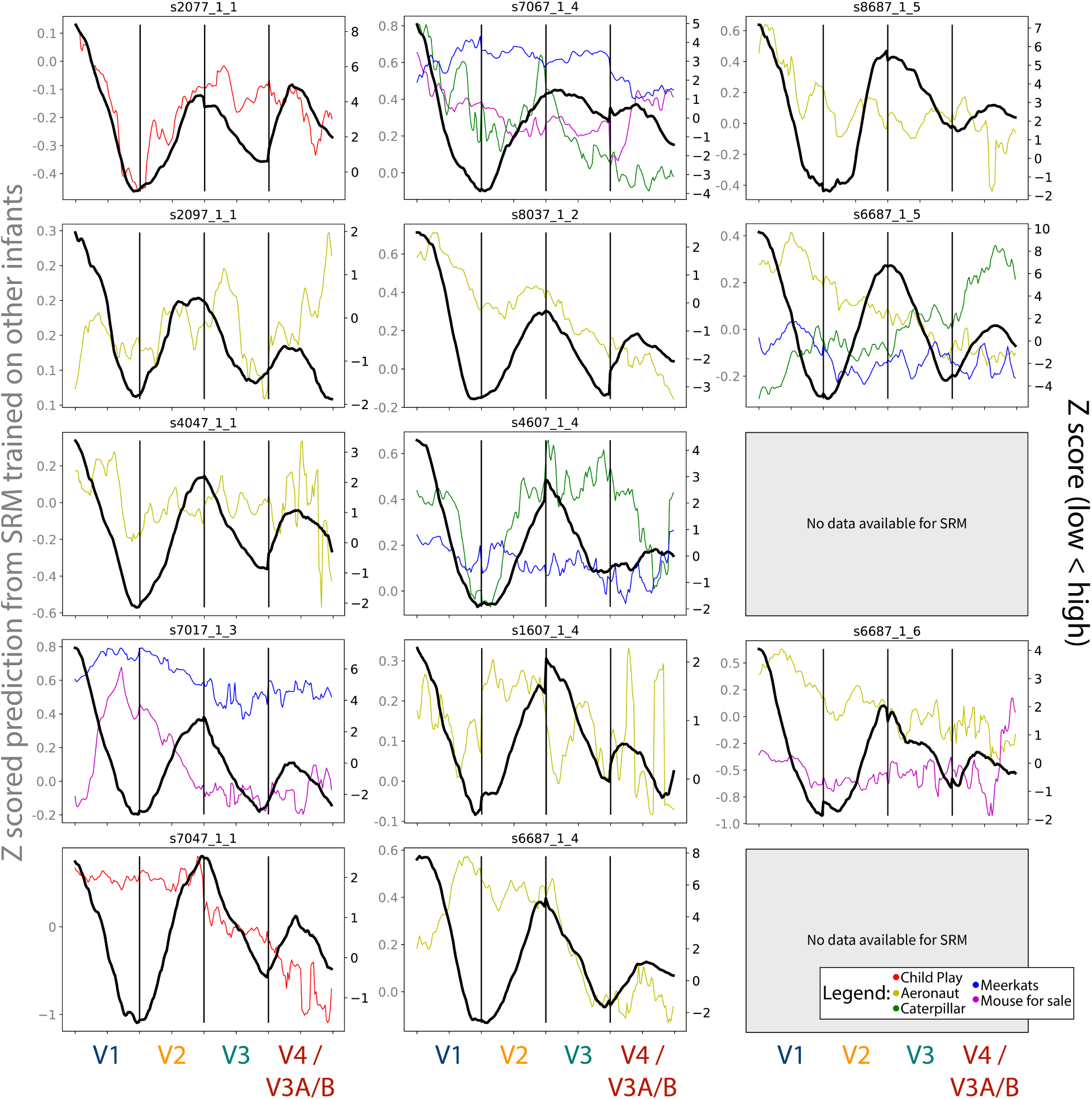
Gradients for the meridian maps predicted using SRM from other infant participants, compared to the task-evoked gradients. The colored lines depict the gradients from each chosen movie that could be used, and their scale is indicated by the Y-axis on the left-hand side. The black line indicates the gradient from the task-evoked map, and their scale is indicated by the Y-axis on the right-hand side. Participants are listed in order of age. Participant data is not reported if the participant did not have SRM-compatible movie data.

**Figure S12:**
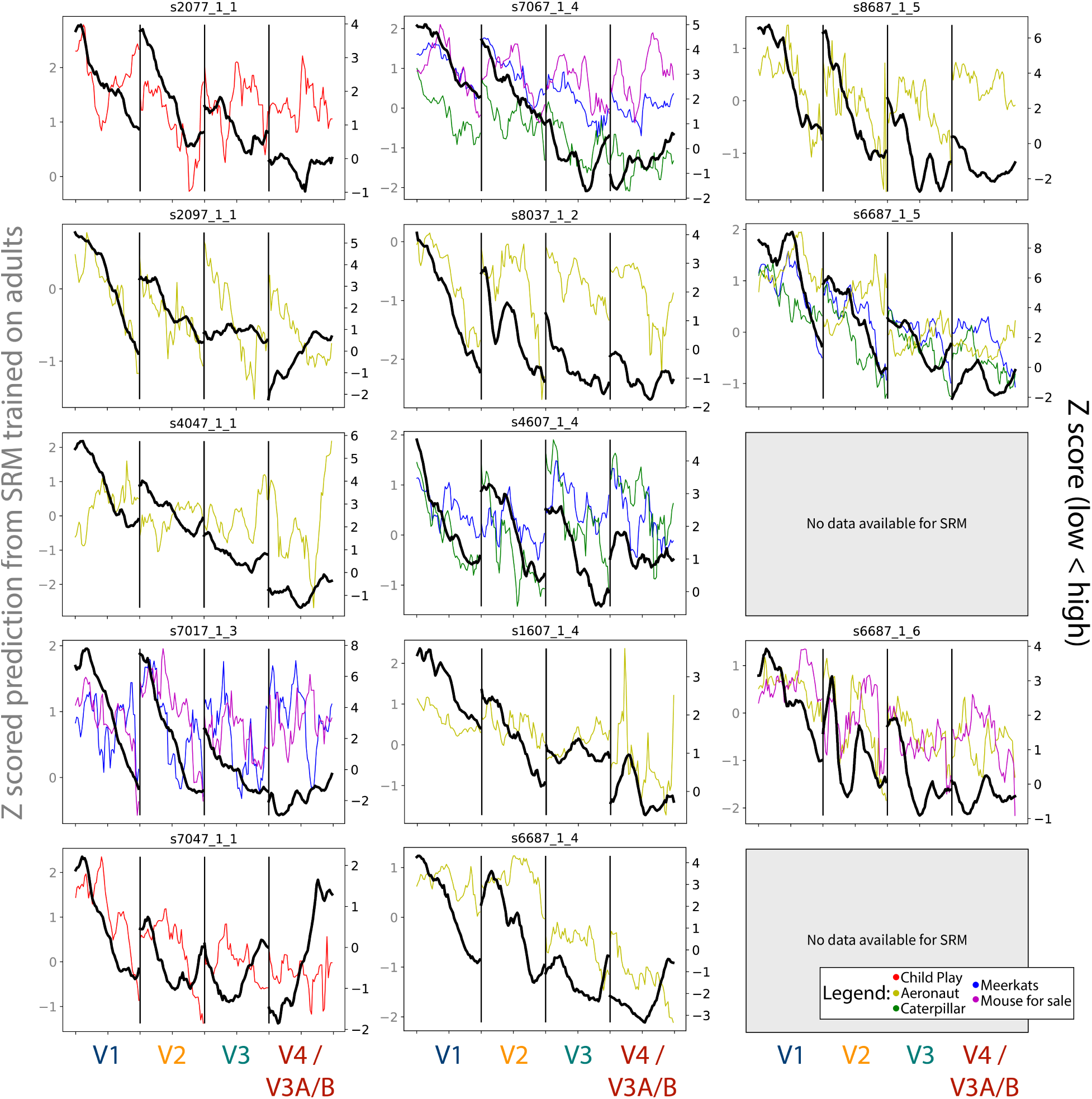
Gradients for the spatial frequency maps predicted using SRM from adult participants, compared to the task-evoked gradients. The colored lines depict the gradients from each chosen movie that could be used, and their scale is indicated by the Y-axis on the left-hand side. The black line indicates the gradient from the task-evoked map, and their scale is indicated by the Y-axis on the right-hand side. Participants are listed in order of age. Participant data is not reported if the participant did not have SRM-compatible movie data.

**Figure S13:**
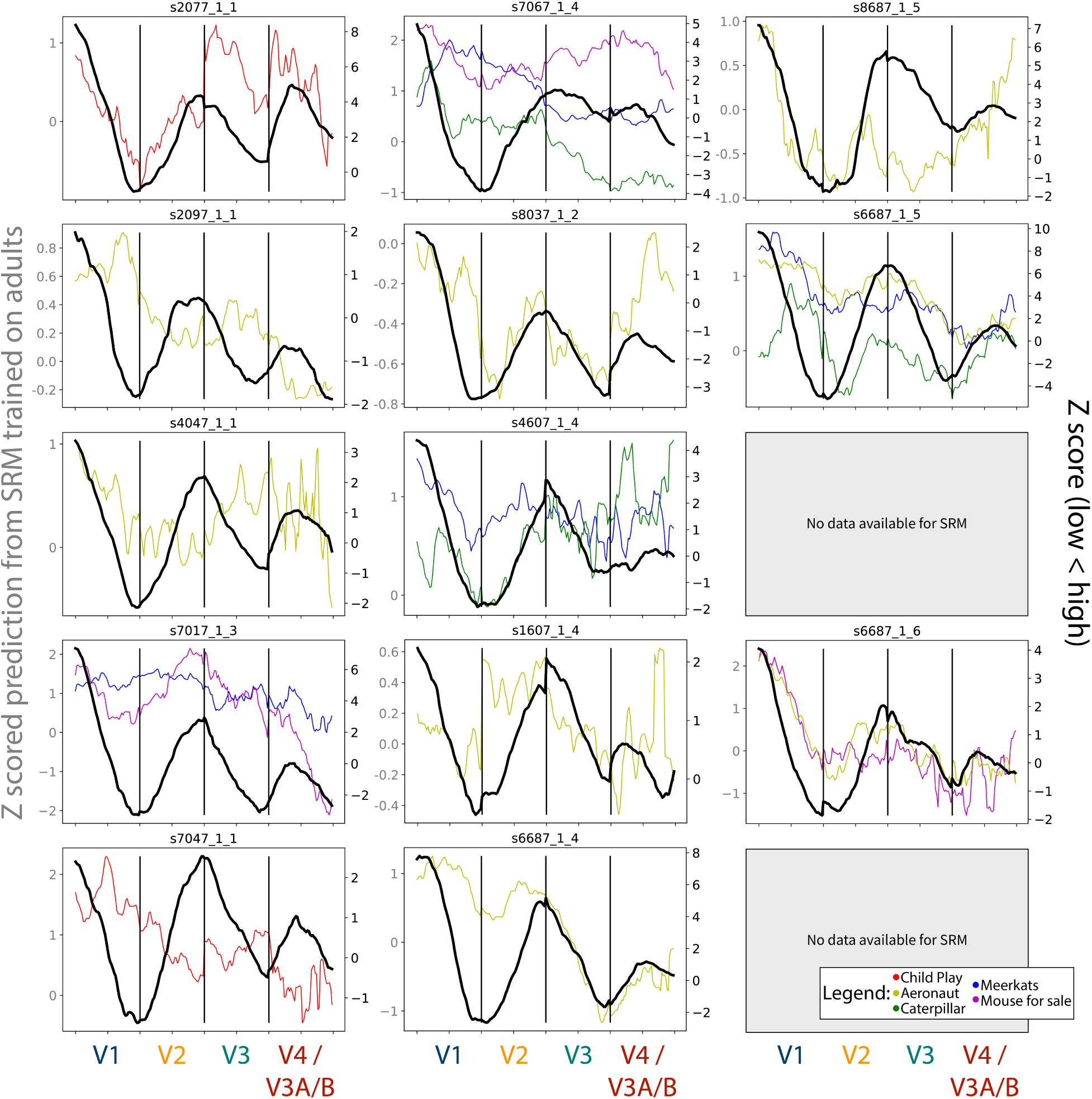
Gradients for the meridian maps predicted using SRM from adult participants, compared to the task-evoked gradients. The colored lines depict the gradients from each chosen movie that could be used, and their scale is indicated by the Y-axis on the left-hand side. The black line indicates the gradient from the task-evoked map, and their scale is indicated by the Y-axis on the right-hand side. Participants are listed in order of age. Participant data is not reported if the participant did not have SRM-compatible movie data.

## Notes

### Competing Interest Statement

The authors have declared no competing interest.

### Summary of Updates

The parameters for adult T1 scans were fixed. Updated some statistic values that changed due to the seed used, aligning the manuscript with the publicly available jupyter notebook. Minor typos were detected and fixed as well.

